# Spatiotemporal control of PIWI compartmentalization by mitochondrial scaffolds defines pachytene piRNA pathway organization

**DOI:** 10.1101/2025.11.02.684826

**Authors:** Xiaoyuan Yan, Chao Wei, Jeffrey M. Mann, Guanyi Shang, Qianyi Wang, Huirong Xie, Elena Y. Demireva, Liangliang Sun, Deqiang Ding, Chen Chen

## Abstract

Pachytene piRNAs are the least understood class of piRNAs in the mammalian male germ line. During meiosis, their biogenesis occurs near mitochondrial outer membrane in germ granules known as intermitochondrial cement (IMC). However, how mitochondrial factors regulate the trafficking of PIWI proteins into and out of the IMC remain poorly understood. Here we show that the cytoplasmic PIWI proteins MILI and MIWI are recruited for pachytene piRNA biogenesis via distinct mitochondrial membrane proteins. Loss of the mitochondrial scaffold protein ASZ1 during meiosis in mice disrupts multiple downstream biogenesis steps, leading to misregulation of MILI and MIWI, failure of IMC formation, and a near-complete loss of mature pachytene piRNAs. Strikingly, despite the drastic depletion of pachytene piRNAs, LINE1 transposon silencing remains unaffected. We identify three classes of pachytene piRNA pathway components that coordinate piRNA production and compartmentalization. Our findings reveal that chromatoid body precursors serve as a central hub for the accumulation of pachytene PIWI-piRNA complexes, thus establishing a connection between IMC-based biogenesis and downstream piRNA function.

## Introduction

PIWI-interacting RNAs (piRNAs) are a class of small noncoding RNAs that associate with PIWI-clad Argonaute proteins to regulate genome stability and germline development^1–3^. Typically 23-35 nucleotides in length, piRNAs guide PIWI proteins to their targets, mediating transposon silencing, RNA degradation through transcriptional and posttranscriptional regulation^4, 5^. piRNA biogenesis involves the processing of long single-stranded precursor RNAs into mature piRNAs through a complex machinery in which several key steps occur in close proximity to mitochondria^2, 6^. In metazoans ranging from flies to mammals, mitochondrial outer membrane serves as a platform for organizing PIWI proteins and piRNA processing enzymes to coordinate piRNA precursor processing albeit precise molecular mechanisms vary^7–11^.

In mammals, pachytene piRNAs represent a unique and highly abundant subclass of piRNAs, expressed specifically during the pachytene stage of meiosis in male germ cells^12–16^. Unlike pre-pachytene piRNAs, which are mainly involved in transposon silencing, pachytene piRNAs have largely unknown targets and functions^17, 18^. Nonetheless, genetic studies have demonstrated that they are essential for spermatogenesis and male fertility^19, 20^. In mice, pachytene piRNAs associate exclusively with two cytoplasmic PIWI proteins MILI and MIWI, most of which do not map to transposable elements, suggesting distinct biological roles beyond genome defense^21–25^.

During pachytene piRNA biogenesis, MILI and MIWI concentrate in the intermitochondrial cement (IMC), germ granules located between clusters of mitochondria in spermatocytes^4, 26^. However, the mechanisms governing how PIWI proteins are recruited to, organized within, and exported from the IMC remain poorly understood. Understanding these mitochondrial-associated processes is critical for elucidating the spatial and temporal regulation of pachytene piRNA production and function.

We have previously shown that mitochondrial protein TDRKH specifically recruits MIWI to the IMC for piRNA processing, while MILI appears to use a distinct, unknown mechanism^27, 28^. In this study, we identify ASZ1 as a mitochondria-anchored scaffold protein that specifically recruits MILI for pachytene piRNA biogenesis. Loss of *Asz1* in postnatal germ cells leads to a near-complete loss of pachytene piRNAs, disrupted IMC structure, and misregulation of piRNA pathway components. We further define distinct classes of pachytene piRNA factors and reveal novel PIWI granules formed post-biogenesis. These findings uncover a critical role for mitochondrial scaffolding in organizing pachytene piRNA production and trafficking during meiosis.

## Results

### ASZ1 selectively recruits MILI, but not MIWI, to mitochondria

Our previous study established that MIWI and MILI enter the IMC through distinct mechanisms during piRNA biogenesis in pachytene spermatocytes^28^. MIWI is initially recruited to the IMC by mitochondria-anchored TDRKH, while how MILI enters the IMC remains unclear. We hypothesize that MILI recruitment is mediated by a different mitochondria-anchored protein. To identify MILI-specific binding proteins, we tested four known mitochondria-anchored piRNA biogenesis factors for their interaction with MILI using a heterologous co-expression system in HeLa cells. ASZ1^29^, TDRKH^30^, GPAT2^31^, and PLD6^11, 32^ all possess transmembrane (TM) regions for mitochondrial membrane anchoring (Fig. 1a). When individually expressed as GFP-tagged proteins in HeLa cells, they colocalized completely with mitotracker, indicating their autonomous anchoring to mitochondria (Fig. 1b). In contrast, FLAG-tagged MILI was diffusely expressed throughout the cytoplasm of HeLa cells (Fig. 1c). When we co-expressed FLAG-MILI with GFP-ASZ1, TDRKH-GFP, GFP-PLD6, or GPAT2-GFP, MILI was specifically recruited to mitochondria by ASZ1, but not by TDRKH, PLD6, or GPAT2, indicating a specific interaction between MILI and ASZ1 (Fig. 1c). By contrast, co-expression of ASZ1 with MIWI did not result in mitochondrial accumulation of MIWI, suggesting that ASZ1 is incapable of recruiting MIWI (Fig. 1d). As a control, MIWI was specifically recruited to mitochondria when co-expressed with TDRKH (Fig. 1e). These results indicate that mitochondria-anchored ASZ1 selectively recruits MILI to mitochondria *in vitro*. We further confirmed the selective MILI-ASZ1 and MIWI-TDRKH interactions *in vivo* using co-immunoprecipitation and Western blotting in mouse testes (Fig. 1f).

**Fig. 1.**
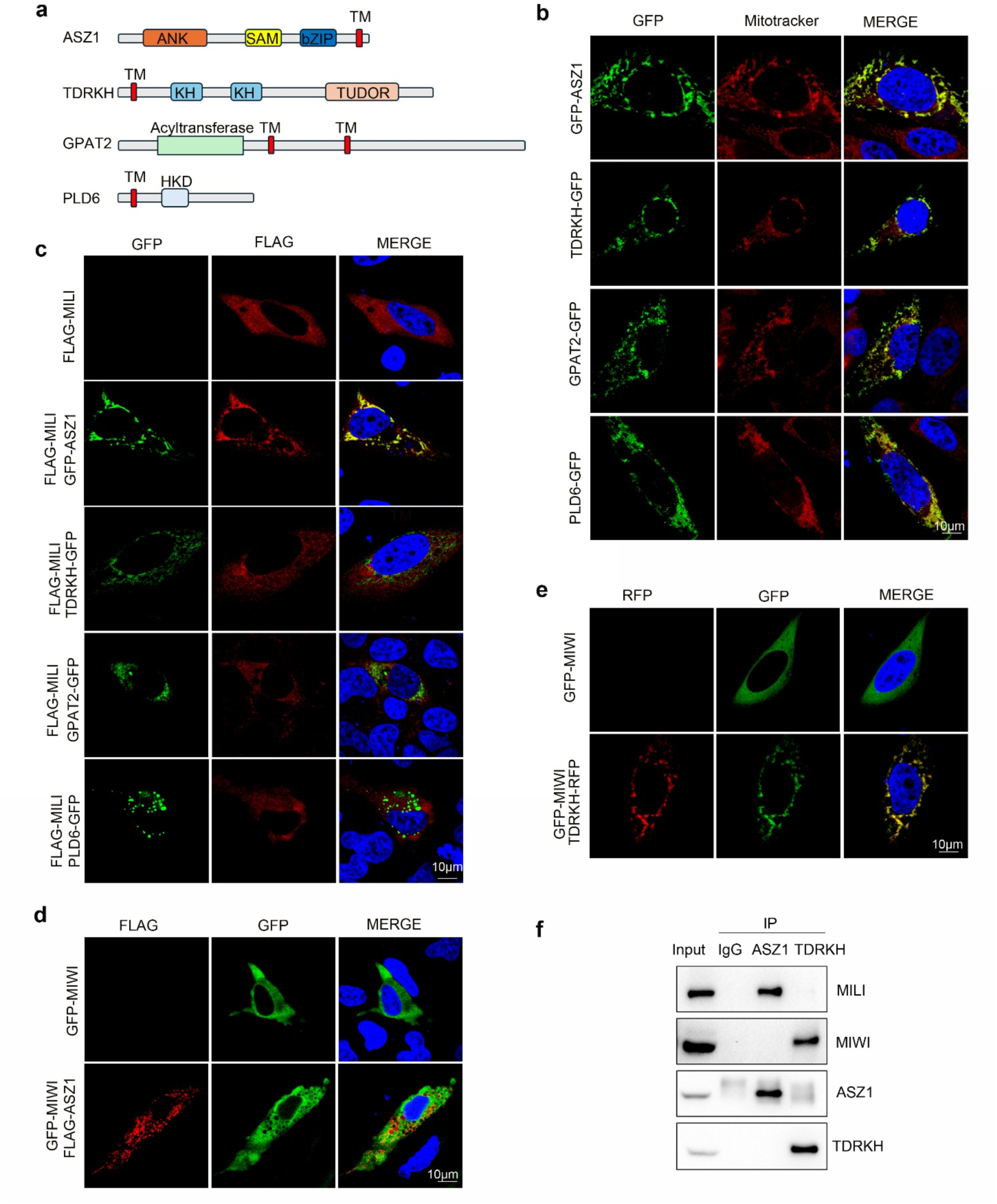
ASZ1 preferentially recruits MILI, but not MIWI, to mitochondria. **a** Domain structures of mitochondrial membrane proteins ASZ1, TDRKH, GPAT2, and PLD6. **b** HeLa cells were transfected with indicated GFP-tagged plasmids. After 24 h, the transfected cells were stained with Mitotracker and then fixed. DNA was stained with DAPI. **c** FLAG-MILI was co-transfected with GFP-ASZ1, TDRKH-GFP, GPAT2-GFP or PLD6-GFP in HeLa cells. The transfected cells were immunostained with anti-FLAG antibody. DNA was stained with DAPI. **d** HeLa cells were transfected with GFP-MIWI, along with FLAG-ASZ1 plasmid. The cells transfected with FLAG-tagged plasmid were immunostained with anti-FLAG antibody. DNA was stained with DAPI. **e** HeLa cells were transfected with GFP-MIWI, along with TDRKH-RFP plasmid. DNA was stained with DAPI. **f** Immunoprecipitation-Western blotting (IP-WB) shows the selective association of ASZ1 with MILI and TDRKH with MIWI in adult WT testes.

### Conditional deletion of *Asz1* in postnatal germ cells disrupts spermiogenesis in mice

We next sought to explore the function of ASZ1 in mouse pachytene piRNA biogenesis. ASZ1 was detected in spermatogonia but was absent in preleptotene spermatocytes. Its expression reappeared in leptotene and zygotene spermatocytes, peaked during mid to late pachytene spermatocytes, and remained detectable in round spermatids (Fig. S1). Previously reported *Asz1* global knockout (KO) mice exhibit defective fetal piRNA biogenesis and spermatogenic arrest at the zygotene stage of meiosis prophase I, precluding the study of its effects on pachytene piRNAs^29^. To circumvent this, we generated mice with conditional deletion of *Asz1* in postnatal male germ cells. This was achieved by first generating an *Asz1* conditional allele (*Asz1^flox^*) with LoxP sites flanking exons 1 and 2 (Fig. 2a, Fig. S2). By combining it with *Stra8*-Cre transgene that enables the deletion of *Asz1* in postnatal day 3 male germ cells, *Stra8*-Cre*, Asz1^flox/-^* conditional KO (referred to as *Asz1^cKO^*) mice were generated, in which *Asz1* is deleted across adult male germ cell lineages (Fig. 2a). Immunofluorescence and Western blotting confirmed the successful ablation of ASZ1 in adult germ cells and testes (Fig. 2b, c). Testes from *Asz1^cKO^* mice were significantly smaller than those from wild-type (WT) controls, weighing approximately 50% of WT testes (Fig. 2d, e). Histological analysis of *Asz1^cKO^*testes revealed spermatogenic arrest at the round spermatid stage, with an absence of elongated spermatids in the seminiferous epithelium (Fig. 2f). This phenotype differs from reported *Asz1^KO^*mice, which exhibited germ cell arrest at the zygotene stage of meiosis, prior to the onset of pachytene piRNAs (Fig. 2f). As a result, only round spermatid-like cells were observed in the epididymides of *Asz1^cKO^*mice (Fig. 2f). Acrosome staining with an anti-ACRV1 antibody further confirmed spermatogenic arrest at the step 1 spermatid stage in *Asz1^cKO^* testes (Fig. 2g). Collectively, these findings demonstrate that ASZ1 expression in postnatal germ cell is crucial for spermiogenesis in mice (Fig. 2h).

**Fig. 2.**
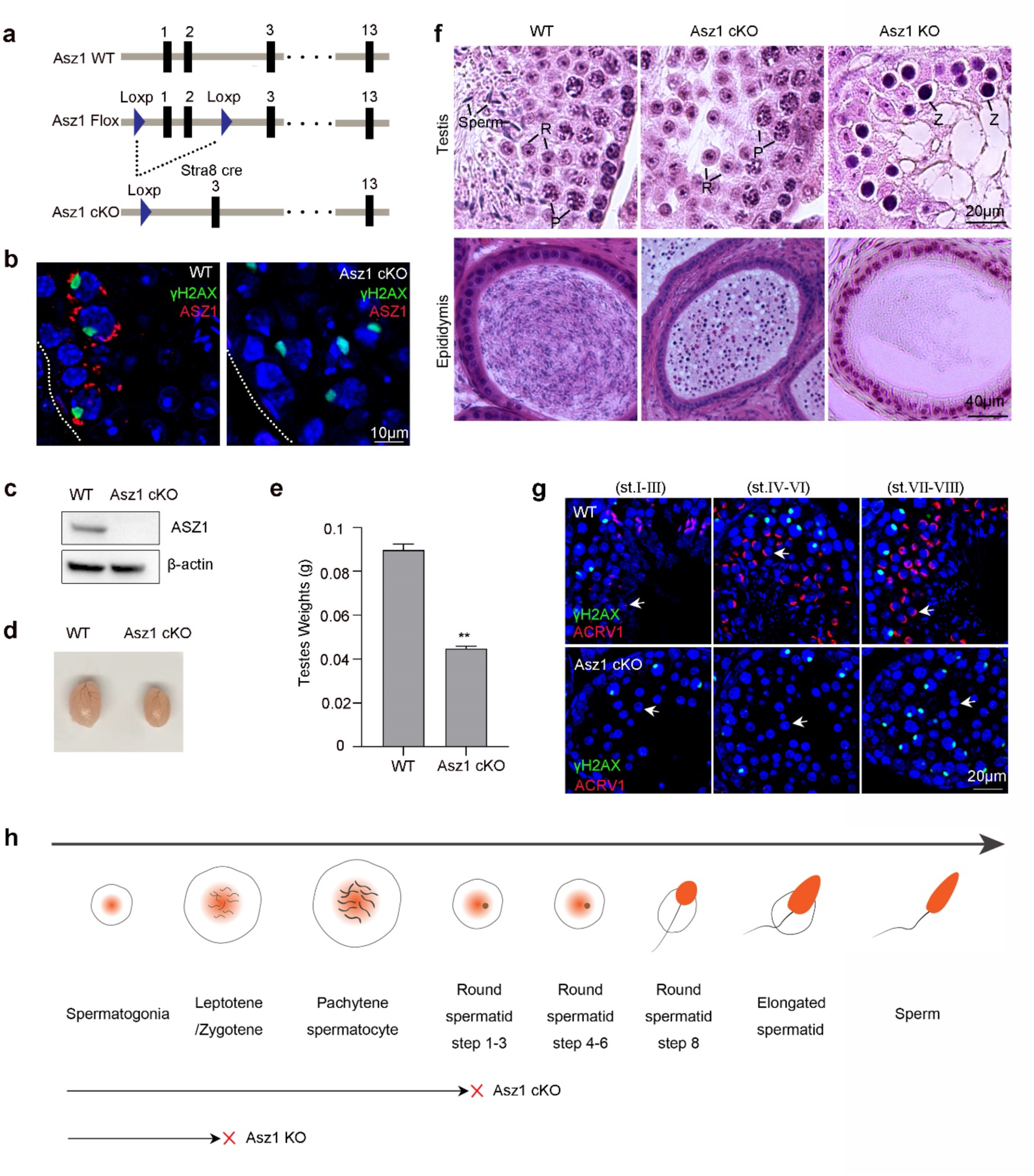
Postnatal germline deletion of *Asz1* causes spermiogenic arrest in mice. **a** Schematic representation of gene targeting strategy for the generation of the *Asz1* conditional allele in mice. *Stra8*-Cre-mediated deletion removes exon 1 and exon 2 of *Asz1*. **b** Immunostaining of ASZ1 in adult WT and *Asz1^cKO^*testes. **c** Western blot analysis of ASZ1 in adult WT and *Asz1^cKO^*testes. **d, e** Testis sizes and weights of WT and *Asz1^cKO^*mice. n = 9. Error bars represent SEM. The P-value was calculated using unpaired t-test. **, P < 0.01. **f** Hematoxylin and eosin staining of testis and epididymis sections from adult WT, *Asz1^cKO^*, and *Asz1^KO^* mice. P pachytene spermatocytes, R round spermatids, Z zygotene spermatocytes. **g** Co-immunostaining of ACRV1 and γH2AX in stage Ⅰ-Ⅲ to stage VII–VIII seminiferous tubules of WT and *Asz1^cKO^* testes. Arrows indicate round spermatids. **h** A schematic diagram illustrating the stages of spermatogenic arrest in *Asz1^cKO^*and *Asz1^KO^* testes.

### ASZ1 loss disrupts MILI localization and causes MIWI aggregation at the IMC

To investigate the impact of postnatal ASZ1 deficiency on the piRNA pathway, we examined the expression levels of key piRNA pathway proteins in *Asz1^cKO^* testes. Western blotting revealed significant reduction in both MILI and MIWI levels, whereas MVH, TDRKH, and Tudor domain proteins (TDRDs) remained comparable to the WT (Fig. 3a). Interestingly, MOV10L1, an RNA helicase involved in pachytene piRNA processing^33^, was also reduced (Fig. 3a). Transmission electron microscopy revealed severe mitochondrial aggregation in *Asz1^cKO^* pachytene spermatocytes (Fig. 3b). Notably, the decreased expression levels of MILI and MIWI, along with clustered mitochondria in *Asz1^cKO^*, closely resembled phenotypes previously reported in *Mov10l1^cKO^* mice^34^.

**Fig. 3.**
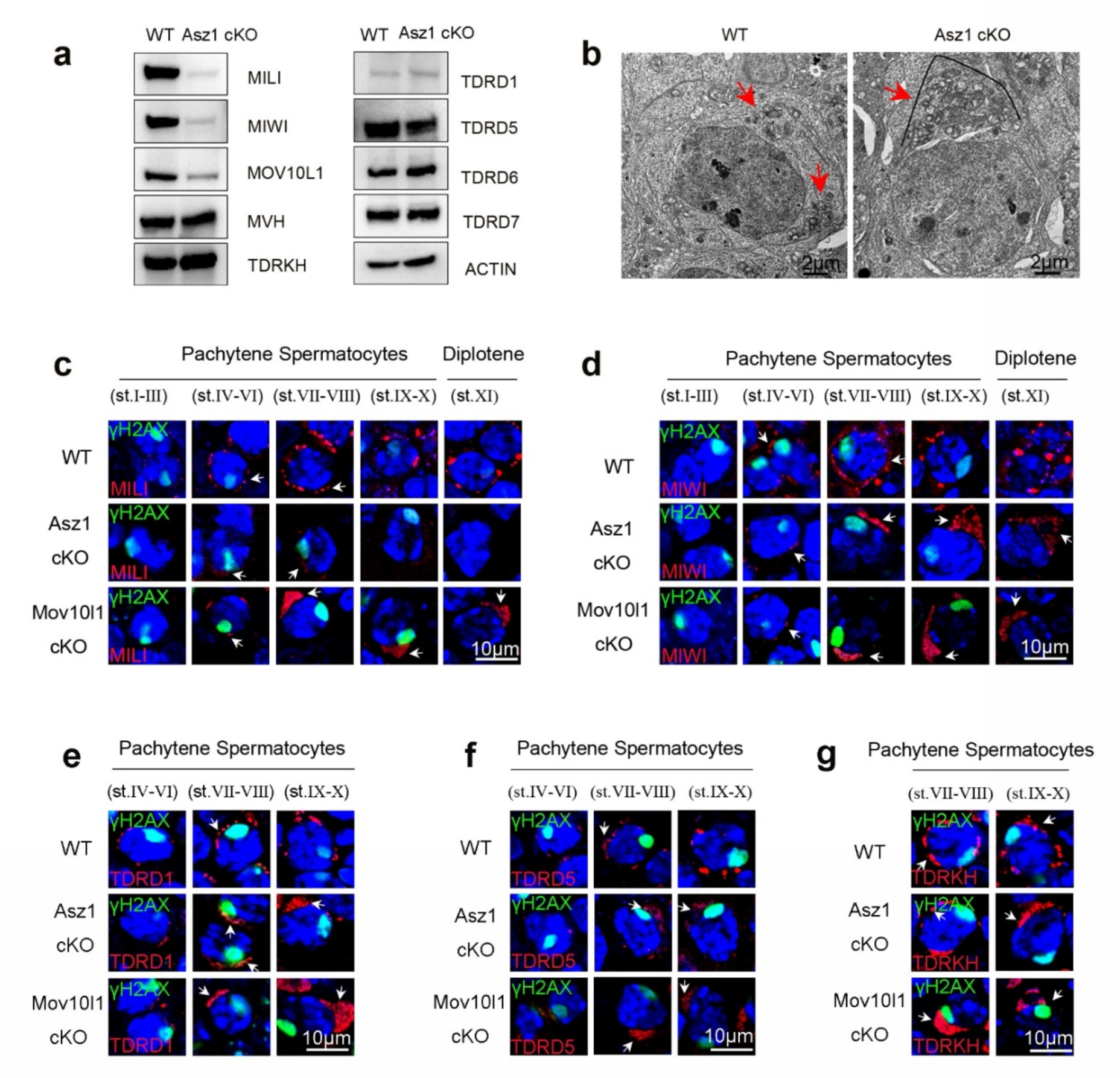
ASZ1 deficiency disrupts MILI localization and causes MIWI aggregation at the IMC. **a** WB of indicated proteins in adult WT and *Asz1^cKO^* testes. **b** Loss of ASZ1 causes the aggregation of mitochondria in spermatocytes. Transmission electron microscopy was performed on pachytene spermatocytes from adult WT and *Asz1^cKO^* testes. Red arrows indicate mitochondria. Black lines indicate mitochondrial aggregation. **c** Immunofluorescence analysis of MILI and γH2AX in adult testis sections from WT, *Asz1^cKO^*, and *Mov10l1^cKO^* mice. White arrows indicate the IMC. **d** Immunofluorescence analysis of MIWI and γH2AX in adult testis sections from WT, *Asz1^cKO^*, and *Mov10l1^cKO^*mice. White arrows indicate the IMC. **e** Immunofluorescence analysis of TDRD1 and γH2AX in adult testis sections from WT, *Asz1^cKO^*, and *Mov10l1^cKO^* mice. White arrows indicate the IMC. **f** Immunofluorescence analysis of TDRD5 and γH2AX in adult testis sections from WT, *Asz1^cKO^*, and *Mov10l1^cKO^*. White arrows indicate the IMC. **g** Immunofluorescence analysis of TDRKH and γH2AX in adult testis sections from WT, *Asz1^cKO^*, and *Mov10l1^cKO^*mice. White arrows indicate the IMC.

Next, we examined the expression and localization of MILI and MIWI in pachytene and diplotene spermatocytes by immunofluorescence. In contrast to abundant expression in the WT spermatocytes, MILI expression was minimal in early-pachytene (Stages I-VI) and mid-pachytene (Stages VII-VIII) and nearly absent in late pachytene (Stages IX-X) and diplotene (Stage XI) spermatocytes (Fig. 3c) in *Asz1^cKO^*testes. This is consistent with our *in vitro* findings that ASZ1 is required for MILI recruitment to mitochondria (Fig. 1). In contrast, in *Mov10l1^cKO^* testes, MILI exhibited an aggregated mitochondrial localization pattern, indicating that the recruitment of MILI to the IMC is not affected by MOV10L1 loss but rather mediated by ASZ1 (Fig. 3c). Consistent with mitochondrial clustering, MIWI displayed an aggregated mitochondrial localization pattern in late pachytene and diplotene spermatocytes of both *Asz1^cKO^* and *Mov10l1^cKO^* testes (Fig. 3d). Similar mitochondrial polarization patterns were observed for TDRD1 and TDRD5 in both *Asz1^cKO^* and *Mov10l1^cKO^* testes (Fig. 3e, f). Notably, ASZ1 deficiency did not affect TDRKH localization to the aggregated mitochondria (Fig. 3g). The specific expression and localization defects of MILI and MIWI are unique, distinguishing from other reported piRNA pathway mutants^25, 34–39^. These findings collectively demonstrate that MILI entry into the IMC is controlled by ASZ1, revealing distinct mechanisms governing MILI and MIWI localization to the IMC by two different mitochondria-anchored proteins, ASZ1 and TDRKH, respectively.

### Postnatal ASZ1 deficiency severely impairs pachytene piRNA biogenesis

We next investigated the role of ASZ1 in postnatal piRNA biogenesis. Radiolabeling of total small RNAs revealed that pachytene piRNAs were barely present in *Asz1^cKO^* testes (Fig. 4a). PIWI immunoprecipitation and RNA labeling experiments further showed that both MILI-bound and MIWI-bound piRNAs were nearly undetectable in *Asz1^cKO^* testes as compared with the WT, confirming that postnatal germline ASZ1 is crucial for pachytene piRNA biogenesis (Fig. 4a). We further sequenced small RNAs isolated from adult WT and *Asz1^cKO^* testes by constructing total small RNA libraries followed by RNA-seq. After normalizing to miRNA counts (21-23nt), MILI-piRNAs were completely absent in *Asz1^cKO^* testes, while MIWI-piRNAs were present at only minimal levels (Fig. 4b). Since the severe reduction in *Asz1^cKO^*pachytene piRNAs resembled that of reported *Mov10l1^cKO^* mice^34^, we compared small RNA profiles between these two mutant mouse lines (Fig. 4b). While total piRNAs were both drastically reduced, however, unlike *Mov10l1^cKO^*in which MILI-piRNAs were slightly retained, MILI-piRNAs were entirely absent in *Asz1^cKO^* testes. This represents the most severely reduced MILI-piRNAs reported to date in mouse models with postnatally deletion of piRNA pathway genes^34–36, 39, 40^, underscoring the specific regulatory role of ASZ1 on MILI function.

**Fig. 4.**
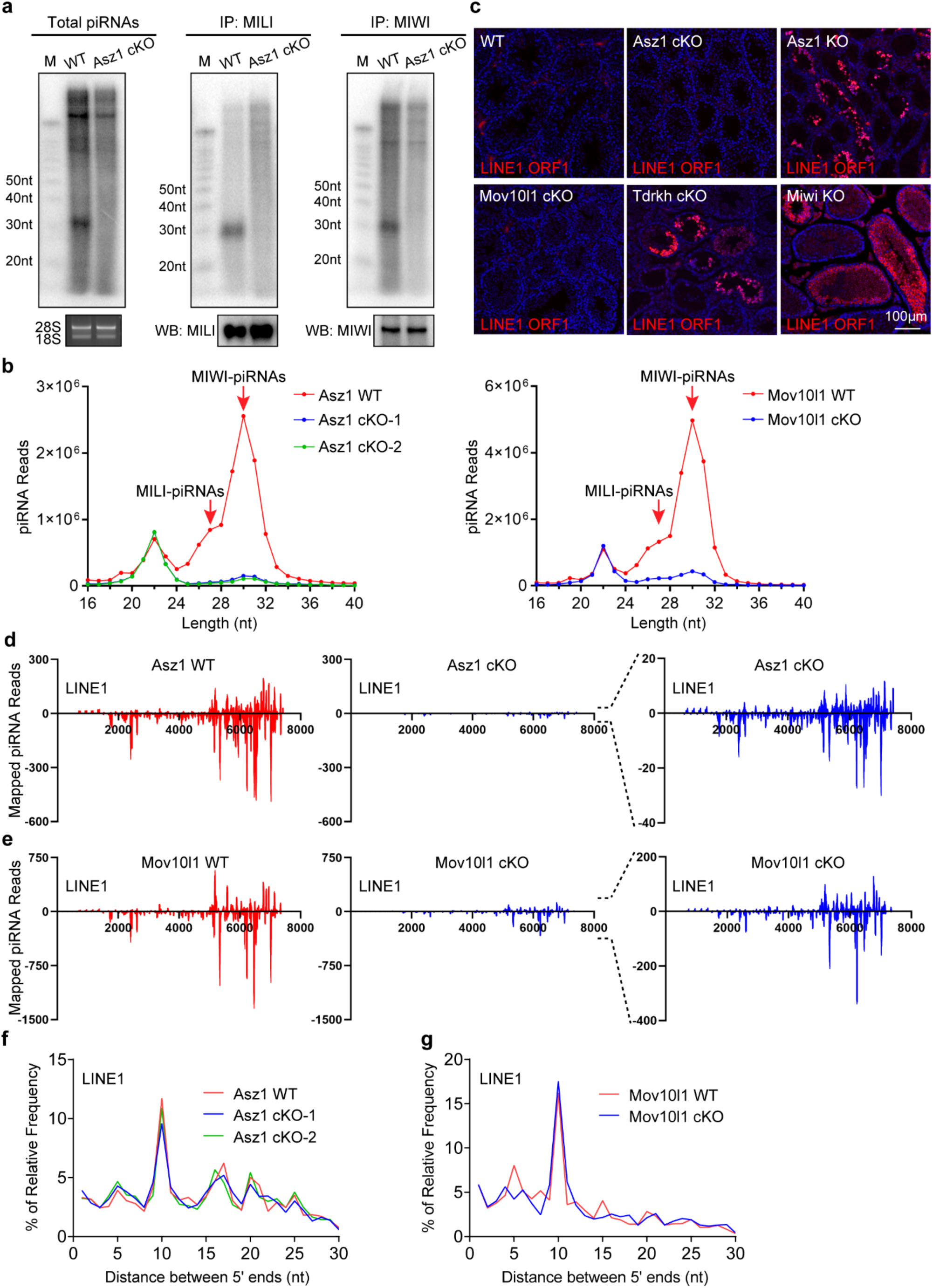
Postnatal deletion of *Asz1* disrupts pachytene piRNA biogenesis without inducing LINE1 de-repression. **a** Severe reduction of piRNAs in *Asz1^cKO^* testes. Total piRNAs, MILI-piRNAs and MIWI-piRNAs were analyzed from adult WT and *Asz1^cKO^* testes. Total RNA was end-labeled with [^32^P]-ATP, separated by 15% TBE urea gel and visualized by autoradiography. 18S and 28S ribosomal RNAs served as loading control. RNA was isolated from immunoprecipitated MILI and MIWI RNPs, end-labeled with [^32^P]-ATP, separated by 15% TBE urea gel and visualized by autoradiography. Western blotting was performed using anti-MILI or MIWI antibodies to show immunoprecipitation efficiency. **b** Length distribution of small RNAs from testicular total small RNA libraries of indicated genotypes. Data were normalized by miRNA reads (21–23 nt). **c** Immunostaining of LINE1 ORF1 in adult WT, *Asz1^cKO^*, *Asz1^KO^*, *Mov10l1^cKO^*, *Tdrkh^cKO^*, and *Miwi^KO^* testes. DNA was stained with DAPI. **d** Graphs display the distribution of piRNAs mapped in the sense and antisense orientations to LINE1 from adult WT (left) and *Asz1^cKO^* (middle) testicular total small RNA libraries. piRNA reads were normalized by miRNA reads (21–23 nt) of each small RNA library. The graph on the right provides a magnified view of the middle graph. Note the differences in the Y-axis scale. **e** Graphs display the distribution of piRNAs mapped in the sense and antisense orientations to LINE1 from adult WT (left) and *Mov10l1^cKO^* (middle) testicular total small RNA libraries. piRNA reads were normalized by miRNA reads (21–23 nt) of each small RNA library. The graph on the right provides a magnified view of the middle graph. Note the differences in the Y-axis scale. **f** The 5′-5′ overlaps between 24-32 nt piRNAs mapped to opposite strands of LINE1 consensus sequence were analyzed using adult WT and *Asz1^cKO^* testicular total small RNA libraries. **g** The 5′-5′ overlaps between 24-32 nt piRNAs mapped to opposite strands of LINE1 consensus sequence were analyzed using adult WT and *Mov10l1^cKO^* testicular total small RNA libraries.

### A minimal level of piRNAs is sufficient to maintain postnatal LINE1 silencing

*Asz1* global KO in mice disrupts fetal piRNA biogenesis and causes significant LINE1 transposon upregulation in germ cells^29^. We next examined the impact of severe reduction of postnatal piRNAs in *Asz1^cKO^* on LINE1 silencing in male germ cells. Surprisingly, despite the severe depletion of piRNAs in *Asz1^cKO^*, LINE1 ORF1 remained suppressed, similar to WT (Fig. 4c). This mimicked the maintenance of LINE1 suppression in *Mov10l1^cKO^*testes, but starkly differing from dramatic upregulation of LINE1 ORF1 in *Asz1^KO^*, *Tdrkh^cKO^*, and *Miwi^KO^* testes in which all MIWI-piRNAs are completely absent (Fig. 4c).

This intriguing observation prompted us to explore the small RNA composition in *Asz1^cKO^* testes in greater detail. Pachytene piRNAs are primarily generated through the PLD6-dependent primary piRNA pathway in the IMC^41, 42^. In *Asz1^cKO^* testes, the residual piRNAs showed reduced proportion of 5′-end U-bias at the first nucleotide and markedly decreased ratio of piRNAs derived from pachytene clusters (Fig. S3), indicating that the primary piRNA pathway is disrupted in *Asz1^cKO^* testes. We then specifically analyzed LINE1-derived piRNAs and found that they were also significantly reduced in *Asz1^cKO^*testes (Fig. 4d). However, the distribution pattern of sense and antisense LINE1-derived piRNAs in *Asz1^cKO^* testes remained comparable to that in the WT (Fig. 4d). This trend was also observed in *Mov10l1^cKO^* testes (Fig. 4e). Additionally, the Ping-Pong signature of the secondary piRNA pathway in LINE1-derived piRNAs was evident and similar between WT and *Asz1^cKO^*, suggesting that the Ping-Pong triggered LINE1 silencing remained functional in *Asz1^cKO^* germ cells (Fig. 4f). Similarly, LINE1-derived piRNAs from *Mov10l1^cKO^* testes showed a comparable Ping-Pong signature (Fig. 4g). Together, these data suggest that LINE1 silencing in postnatal germ cells can be maintained with a minimal level of piRNAs and the vast majority of LINE1-independent pachytene piRNAs are essential for spermiogenesis.

### ASZ1 recruits MOV10L1 to the IMC

To interrogate how ASZ1 regulates pachytene piRNA biogenesis, we performed ASZ1 immunoprecipitation and mass spectrometry (IP-MS) in adult WT testes to identify ASZ1-interacting proteins (Fig. 5a). MOV10L1 and MILI were among the top interactors identified, and mitochondrial membrane protein GPAT2 was also significantly enriched in the ASZ1 complex (Fig. 5a). Mitochondria-anchored TDRKH, however, was not detectable. Notably, MIWI and MVH were not enriched in ASZ1-IP (Fig. 5a). Further co-immunoprecipitation and Western blotting confirmed that ASZ1 interacts with MOV10L1 and MILI, but not MIWI or MVH in adult testes (Fig. 5b).

**Fig. 5.**
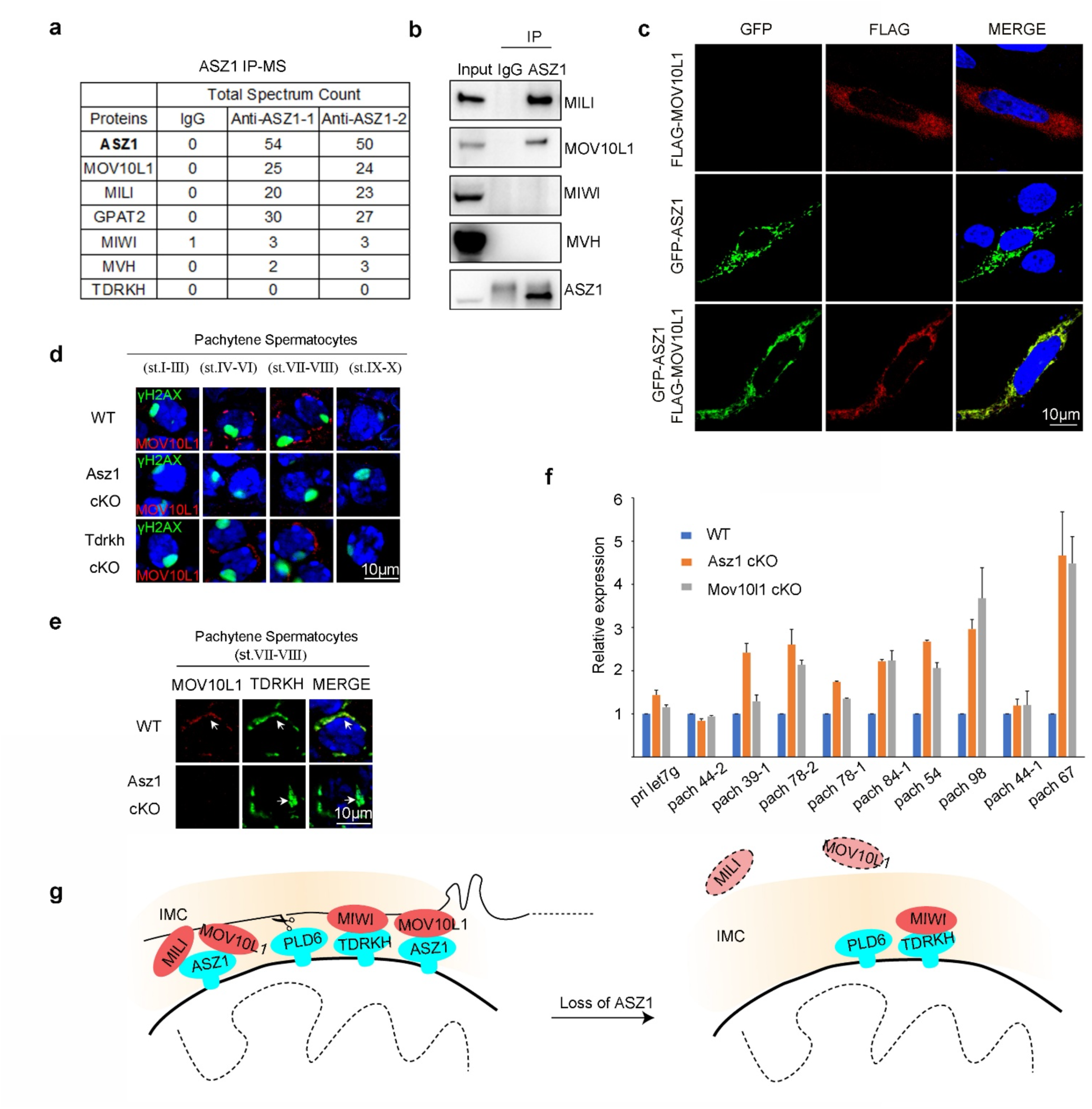
ASZ1 interacts with MOV10L1 to promote pachytene piRNA biogenesis. **a** Identification of ASZ1-interacting proteins in adult WT testes by IP and mass spectrometry. The total spectral counts for the indicated proteins are shown. **b** IP-WB analysis showing the association of ASZ1 with MILI and MOV10L1 in adult WT testes. **c** HeLa cells were transfected with indicated GFP-tagged and/or FLAG-tagged plasmids. Cells transfected with FLAG-tagged plasmid were immunostained with anti-FLAG antibody. DNA was stained with DAPI. **d** Immunofluorescence of MOV10L1 and γH2AX in adult WT, *Asz1^cKO^*, and *Tdrkh^cKO^*testes. **e** Co-immunostaining of MOV10L1 and TDRKH in adult WT, *Asz1^cKO^*, and *Tdrkh^cKO^* testes. White arrows indicate the IMC. **f** piRNA precursor transcript levels were quantified by RT-qPCR in WT, *Asz1^cKO^*, and *Mov10l1^cKO^* testes. miRNA precursor pri-let7g served as a control. n = 3; error bars represent SEM. **g** A proposed model for the role of ASZ1 in pachytene piRNA biogenesis. Loss of ASZ1 profoundly impairs the biogenesis of both MILI- and MIWI-bound piRNAs, attributable to defective localization of MILI and MOV10L1 to the IMC.

Since MOV10L1 is a major ASZ1 interactor and an RNA helicase bound to piRNA precursors required for pachytene piRNA biogenesis, we next tested whether mitochondria-anchored ASZ1 could directly recruit MOV10L1 to mitochondria. When FLAG-tagged MOV10L1 was expressed in HeLa cells, it exhibited a diffuse cytoplasmic distribution (Fig. 5c). When co-expressed with ASZ1, however, MOV10L1 was significantly recruited to colocalize with ASZ1 on mitochondria, suggesting the ability of ASZ1 to directly recruit MOV10L1 for piRNA processing (Fig. 5c).

We then examined the effect of ASZ1 deficiency on MOV10L1 expression and localization during pachytene piRNA biogenesis in mice. In WT testes, MOV10L1 was expressed in stage IV-VIII pachytene spermatocytes and colocalized with mitochondria-anchored TDRKH (Fig. 5d, e). However, MOV10L1 was largely undetectable in *Asz1^cKO^* pachytene spermatocytes, suggesting that ASZ1 is required for proper MOV10L1 localization (Fig. 5d, e). By contrast, MOV10L1 localization was comparable to WT in *Tdrkh^cKO^* pachytene spermatocytes, suggesting that ASZ1 but not TDRKH selectively regulates MOV10L1 (Fig. 5d). Consistent with the reported accumulation of pachytene piRNA precursors in *Mov10l1^cKO^* testes^34^, we observed increased levels of pachytene piRNA precursors in *Asz1^cKO^* testes (Fig. 5f). Taken together, these findings indicate that ASZ1 acts as a mitochondria-anchored adaptor protein that recruits MILI and MOV10L1 to the IMC to engage pachytene piRNA biogenesis (Fig. 5g).

### Distinct PIWI granules develop away from the IMC in late spermatocytes

The IMC is defined as electron dense germ granules observed in-between mitochondria in germ cells, yet its relationship with PIWI proteins remains loosely defined. Since ASZ1 is immobilized on the mitochondrial surface to promote piRNA production, we used it as a mitochondrial marker to track PIWI protein localization dynamics during pachytene piRNA biogenesis. In WT testes, when co-stained with ASZ1, both MILI and MIWI showed high colocalization (∼70%) with ASZ1 in stage VII-VIII pachytene spermatocytes (Fig. 6a, d). When progressing to stage IX-X, colocalization ratio of ASZ1 with MILI and MIWI decreased, with the noticeable emergence of several newly formed PIWI-alone granules adjacent to ASZ1 positive foci (Fig. 6a, d). Strikingly, in stage XI diplotene spermatocytes, these new PIWI granules become more prominent and completely segregated from ASZ1 foci (Fig. 6a, d). As a result, these large PIWI granules showed only little colocalization (<20%) with ASZ1 (Fig. 6a, d). The de novo formation of these new PIWI granules away from the IMC was further confirmed by colocalization of MILI and MIWI with another mitochondria-anchored protein TDRKH (Fig. 6b, e), suggesting the dynamic translocation of PIWI proteins following processing from the IMC to new subcellular compartments.

**Fig. 6.**
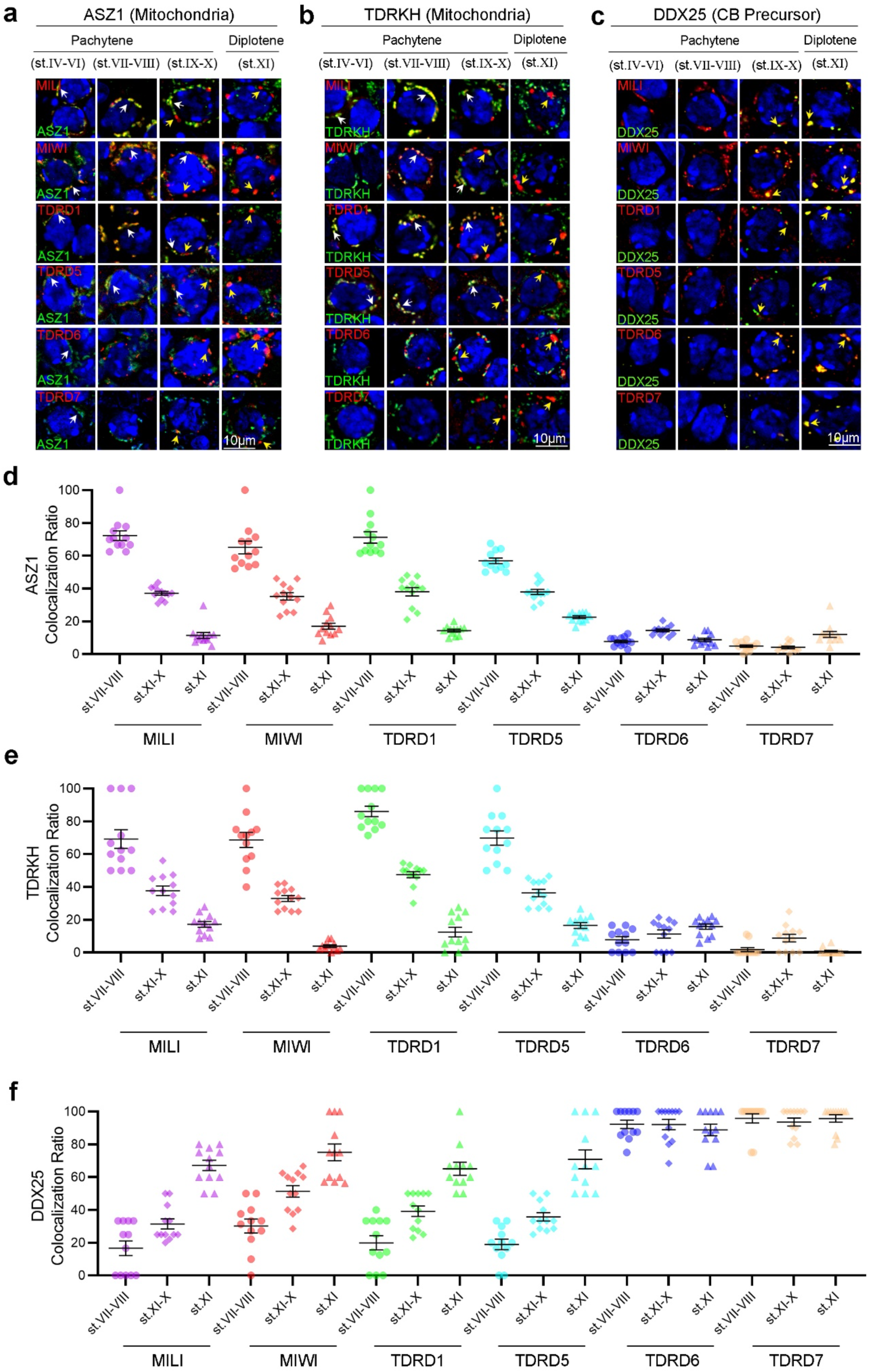
Spatiotemporal dynamics of PIWI granule formation during meiosis. **a-c** Co-immunostaining of indicated piRNA pathway proteins with ASZ1, TDRKH, or DDX25 in adult WT testis sections. White arrows indicate the IMC. Yellow arrows indicate CB precursors. **d** Quantification of the colocalization ratio between indicated piRNA pathway proteins and ASZ1 in pachytene and diplotene spermatocytes. n = 12 spermatocytes from three mice, data are presented as mean ± SEM. **e** Quantification of the colocalization ratio between indicated piRNA pathway proteins and TDRKH in pachytene and diplotene spermatocytes. n = 12 spermatocytes from three mice, data are presented as mean ± SEM. **f** Quantification of the colocalization ratio between indicated piRNA pathway proteins and DDX25 in pachytene and diplotene spermatocytes. n = 12 spermatocytes from three mice, data are presented as mean ± SEM.

TDRD1 and TDRD5 are known IMC enriched proteins crucial for piRNA biogenesis^36, 43, 44^. Like MILI and MIWI, their colocalization with ASZ1 or TDRKH decreased during spermatocyte progression from pachytene to diplotene (Fig. 6a, b, d, e). The dissociation of MILI, MIWI, TDRD1, and TDRD5 from mitochondrial ASZ1 and TDRKH marked IMC strongly suggests their transition together with PIWI proteins to form new PIWI granules. This coincides with the completion of piRNA processing and maturation of piRNA-loaded PIWI proteins exiting the IMC for downstream physiological function.

### Newly emerged PIWI granules merge with TDRD6 and TDRD7 in CB precursors

To trace and characterize the emergence of newly formed PIWI granules post-stage VIII, we analyzed TDRD6 and TDRD7, late-expressed Tudor proteins implicated in piRNA pathway function but not directly involved in piRNA biogenesis^37, 45, 46^. TDRD6 and TDRD7 started to express in stage VII-VIII pachytene spermatocytes and peaked in diplotene spermatocytes as large granules (Fig. 6a-c and Fig. S4). Notably, they had minimal colocalization with mitochondria-anchored ASZ1 or TDRKH in the IMC throughout meiotic progression (Fig. 6a, b, d, e). TDRD6 and TDRD7 instead showed nearly complete overlap in expression timing and colocalization with DDX25, a well-established marker of CB precursors^47^ (Fig. 6c, f). In contrast, MIWI, MILI, TDRD1, and TDRD5 showed a distinct colocalization pattern with DDX25, with their colocalization ratios rising from approximately 20% at stage VII–VIII to about 70% by stage XI of meiotic spermatocytes (Fig. 6c, f). This shift coincides with the reduction in their colocalization with mitochondrial ASZ1 or TDRKH, signifying translocation of PIWI proteins from the IMC to CB precursors.

### Three classes of pachytene piRNA factors reveal dynamic PIWI translocation between germ granules

Based on temporospatial localization of piRNA factors shown above, we classify piRNA pathway proteins into three categories during pachytene piRNA biogenesis: (1) Mitochondria-associated factors (e.g., ASZ1, TDRKH), mitochondrial-anchored proteins restricted to the outer mitochondrial membrane throughout meiosis; (2) CB precursor-specific factors (e.g., TDRD6, TDRD7), expressed in mid-to-late pachytene, localized exclusively to DDX25-positive CB precursors with no IMC overlap; and (3) transitional factors (e.g., MILI, MIWI, TDRD1, and TDRD5), which initially colocalize with mitochondria-associated factors but later translocate to CB precursors. Collectively, these three classes of piRNA factors reflect the spatiotemporal dynamics of piRNA biogenesis during meiotic prophase I, suggesting a directional flow of PIWI-piRNA complexes from the IMC to CB precursors for germ granule reorganization.

### ASZ1 loss impedes the translocation of transitional piRNA factors from the IMC to CB precursors

We next examined the impact of mitochondrial factor loss on the localization of transitional factors during pachytene piRNA biogenesis. In *Asz1^cKO^* spermatocytes, MILI expression was markedly reduced (Fig. 7a). Co-staining with the mitochondrial TDRKH revealed mitochondrial aggregation and a loss of MILI in stage X spermatocytes (Fig. 7a). CB precursors marked by DDX25 appeared smaller than those in WT at this stage (Fig. 7a). Notably, MIWI, TDRD1, and TDRD5 were aberrantly retained around aggregated mitochondria in *Asz1^cKO^*cells (Fig. 7b-d), in contrast to WT, where PIWI granules emerge away from the IMC and become prominent at stage X (Fig. 7b-d). Consist with this, colocalization between transitional factors and the CB precursor marker DDX25 was minimal in *Asz1^cKO^* stage X spermatocytes, in contrast to the strong overlap observed in WT (Fig. 7a-d). This suggests a failure in the translocation of transitional factors from the IMC to CB precursors upon ASZ1 loss. Accordingly, germ granules containing the late-expressed CB precursor-specific piRNA factors TDRD6 and TDRD7 remained spatially separated from the IMC in stage X *Asz1^cKO^* spermatocytes, despite a noticeable reduction in size, possibly due to the failure to receive transitional factors from the IMC (Fig. 7e, f). Despite this, their colocalization with the CB precursor marker DDX25 was maintained, indicating that the incorporation of TDRD6 and TDRD7 into CB precursors occurs independently of the translocation of transitional piRNA factors (Fig. 7e, f). Similarly, *Mov10l1^cKO^* spermatocytes showed IMC aggregation, retention of transitional piRNA factors at the IMC, and reduced CB precursor size during the transition from pachytene to diplotene (Fig. S5). Together, these findings highlight how disruption of pachytene piRNA biogenesis impairs the translocation of piRNA factors between germ granules, emphasizing the critical role of their dynamic trafficking through distinct subcellular compartments during meiotic prophase I.

**Fig. 7.**
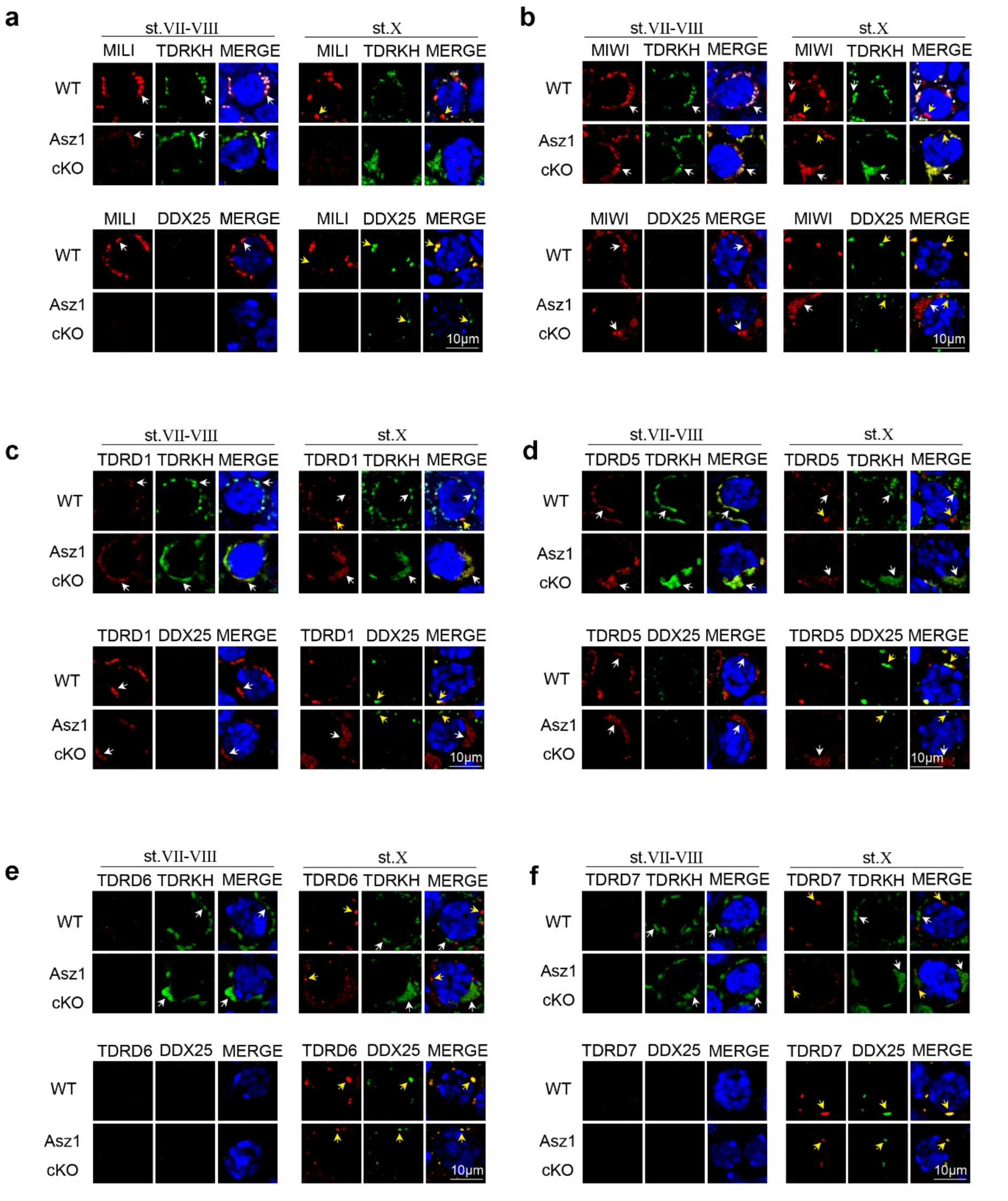
ASZ1 deficiency disrupts translocation of piRNA pathway proteins from the IMC to CB precursors in pachytene spermatocytes. **a** Co-immunostaining of MILI with TDRKH or DDX25 in adult testis sections from WT and *Asz1^cKO^* mice. **b** Co-immunostaining of MIWI with TDRKH or DDX25 in adult testis sections from WT and *Asz1^cKO^* mice. **c** Co-immunostaining of TDRD1 with TDRKH or DDX25 in adult testis sections from WT and *Asz1^cKO^* KO mice. **d** Co-immunostaining of TDRD5 with TDRKH or DDX25 in adult testis sections from WT and *Asz1^cKO^*mice. **e** Co-immunostaining of TDRD6 with TDRKH or DDX25 in adult testis sections from WT and *Asz1^cKO^* mice. **f** Co-immunostaining of TDRD7 with TDRKH or DDX25 in adult testis sections from WT and *Asz1^cKO^* mice. White arrows indicate the IMC. Yellow arrows indicate CB precursors.

### Defective PIWI translocation results in CB malformation in round spermatids

To further investigate the impact of impaired translocation of transitional piRNA factors from the IMC to CB precursors, we examined CB formation in *Asz1^cKO^*round spermatids.

Transmission electron microscopy revealed that CBs in *Asz1^cKO^*spermatids were less electron-dense and appeared fragmented compared to those in WT (Fig. 8a). As a result, transitional factors MILI, MIWI, TDRD1, and TDRD5 were largely absent from *Asz1^cKO^* CBs (Fig. 8b). In contrast, the CB precursor-specific factors TDRD6 and TDRD7 remained detectable in the malformed CBs, confirming that their incorporation into the CB occurs independently of the IMC, transitional factors, and piRNA biogenesis (Fig. 8c).

**Fig. 8.**
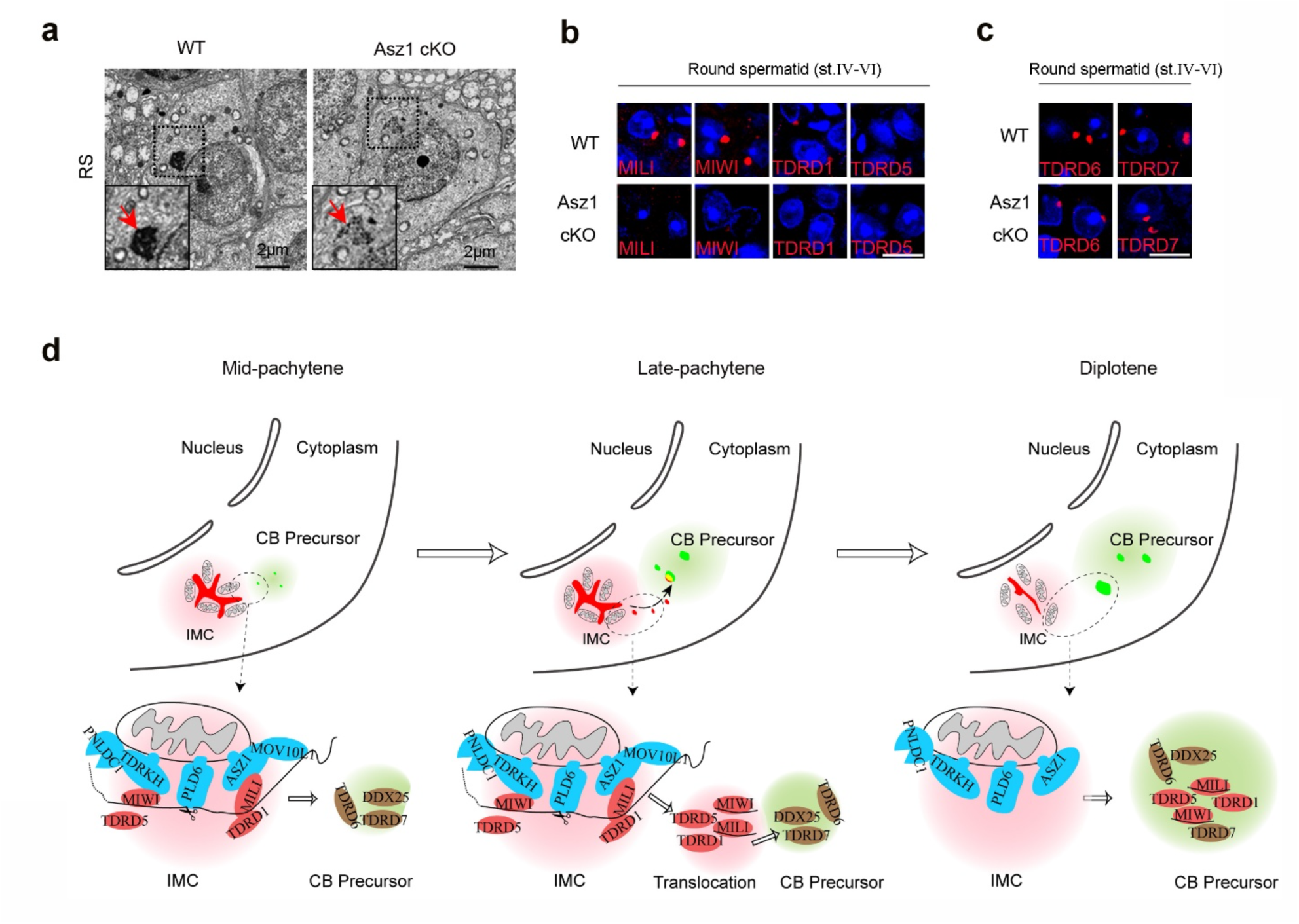
Loss of ASZ1 leads to chromatoid body fragmentation and absence of transitional piRNA factors in round spermatids. **a** ASZ1 deficiency causes chromatoid body fragmentation in round spermatids. Transmission electron microscopy of round spermatids from adult WT and *Asz1^cKO^* testes. Chromatoid bodies are enlarged in the insets and indicated by red arrows. **b** Immunostaining of MILI, MIWI, TDRD1 and TDRD5 in round spermatids from adult WT and *Asz1* cKO testes, scale bar, 10µm. **c** Immunostaining of TDRD6 and TDRD7 in round spermatids from adult WT and *Asz1^cKO^* mice, scale bar, 10µm. **d** A proposed model of stage-specific coordination of pachytene piRNA biogenesis and PIWI compartmentalization during meiosis.

## Discussion

Pachytene piRNAs remain poorly understood, with elusive molecular targets. By examining their biogenesis during mouse meiotic prophase I, we show that two PIWI proteins enter pachytene piRNA biogenesis through distinct mechanisms. We further define the spatiotemporal dynamics of piRNA factors at the IMC and the functional organization of PIWI granules, illuminating the fate of pachytene piRNAs after biogenesis.

Despite MILI and MIWI associate with nearly identical sets of pachytene piRNAs, we establish a model in which two distinct mitochondrial membrane proteins control the entry of MILI and MIWI into pachytene piRNA biogenesis during meiosis. We previously discovered that mitochondrial anchored TDRKH specifically recruits MIWI, but not MILI, to the IMC^28^. Here, we identified ASZ1 as a mitochondrial membrane protein that specifically interacts with MILI. Loss of ASZ1 results in MILI destabilization and a near-complete depletion of pachytene piRNAs. In contrast, MIWI localization to mitochondria remains unaffected, likely due to the continued presence of TDRKH. In addition to MILI, ASZ1 was found to interact with MOV10L1 in the mouse testis. Notably, ASZ1 deficiency compromises both the recruitment and steady-state levels of MOV10L1. Given that MOV10L1 directly binds piRNA precursors^33^, its reduction likely underlies the drastic loss of pachytene piRNAs observed in *Asz1^cKO^* testes, potentially due to the unavailability of piRNA precursors. Consequently, MIWI-piRNA production also fails, not because MIWI recruitment is impaired, but because piRNA precursors are inaccessible. The differential recruitment of MILI and MIWI may potentiate a divergence in the path of MILI and MIWI take to separate their functional roles (e.g. MIWI’s role in post-meiotic silencing vs MILI’s pre-meiotic roles). Indeed, two recent reports showed ASZ1-MILI interaction is conserved for pre-pachytene piRNA production in mouse fetal germ cells^48, 49^. In flies, mitochondrial ASZ1 dimerizes with Daedalus to recruit Piwi for piRNA processing^8, 9^. Therefore, despite the common theme of mitochondrial proteins recruiting of PIWI proteins for piRNA biogenesis, the molecular mechanisms underlying PIWI recruitment vary across species.

One striking finding is that LINE1 silencing remains intact in *Asz1^cKO^*testes, even though these mice exhibit the most severe reduction in pachytene piRNAs among known piRNA-deficient models with spermiogenic arrest^25, 28, 34–36, 38–40, 50^. Postnatal ablation of ASZ1 eliminates MILI-piRNAs and nearly abolishes MIWI-piRNAs, yet LINE1 remain repressed, similar to *Mov10l1^cKO^*testes that retain both trace MILI- and MIWI-piRNAs^33^. In contrast, *Miwi^KO^* and *Tdrkh^cKO^* mice, which completely lack MIWI-piRNAs, display LINE1 activation^25, 28^. These findings indicate that defective LINE1 silencing is not the primary cause of spermiogenic arrest in *Asz1^cKO^* testes. Instead, they support that even minimal levels of transposon-targeting piRNAs are sufficient to preserve genome integrity during meiosis. Indeed, we observed residual LINE1 piRNAs and normal ping-pong signature in *Asz1^cKO^* mice. Therefore, the vast majority of pachytene piRNAs, unrelated to transposon silencing, likely fulfill essential functions in germ cell development, possibly through post-transcriptional regulation or gene silencing mechanisms that remain to be elucidated^51–54^.

Our study uncovers unprecedented insights into the temporospatial regulation of PIWI granule dynamics during pachytene piRNA biogenesis. Using mitochondrial proteins ASZ1 and TDRKH as spatial references, we delineate the coordinated actions of three classes of piRNA biogenesis factors during IMC assembly and disassembly: mitochondria-associated factors, transitional factors, and CB precursor–specific factors (Fig. 8c). ASZ1 and TDRKH act as mitochondrial scaffolds that recruit PIWI proteins, ensuring precursor processing, loading, and trimming occur in close mitochondrial proximity. Upon piRNA maturation, MIWI and MILI translocate from the IMC to the CB precursor together with transitional factors TDRD1 and TDRD5. PIWI granule dissociation from the IMC is likely mediated by TDRD1 through phase separation, as it can tether PIWI proteins into biomolecular condensates^35, 48^. TDRD5, which binds piRNA precursors essential for processing^36^, may also facilitate this transition. Our data further indicate that TDRD1 and TDRD5 converge with newly expressed Tudor proteins TDRD6 and TDRD7 to form CB precursors—the principal destination of PIWI granules after IMC departure. The onset of TDRD6/7 expression from stage VII–VIII and onward coincides with peak piRNA biogenesis, preparing for IMC exit. Notably, TDRD6/7 recruitment to CB precursors occurs independently of piRNA biogenesis, as ASZ1 deletion does not affect their localization. However, without PIWI–piRNA complexes, CB precursors fail to mature properly, leading to malformed CBs in round spermatids. This highlights the importance of IMC components merging with CB precursors as a crucial step for CB formation and function.

In summary, mitochondrial surface proteins coordinate with transitional piRNA factors to initiate pachytene piRNA biogenesis and IMC formation during meiosis. Sequential Tudor protein engagement then drives the emergence of new PIWI granules away from the IMC, establishing temporospatial order from biogenesis to functional compartmentalization. The CB precursor arises as a key site for pachytene piRNA activity, whose precise role in spermatogenesis, beyond transposon silencing, warrants further investigation.

## Methods

### Ethics statement

All animal procedures were approved by the Institutional Animal Care and Use Committee of Michigan State University. All experiments with mice were performed ethically in accordance with the Guide for the Care and Use of Laboratory Animals and institutional guidelines.

### Mouse strains

For the generation of *Asz1^flox^* mice by CRISPR-Cas9 genome editing, two synthetic single guide (sg) RNAs (Synthego) were used to make RNPs which were introduced along with a homology directed repair (HDR) template into C57BL/6N zygotes via pronuclear microinjection. Protospacer (N)20 and PAM sequences corresponding to sgRNAs were 5’- TGTCACGAACGAGTACCACA -GGG-3’ at 404bp before the start *Asz1* exon 1, and 5’- ACTGAGCATGGTGATAGTTG -GGG-3’ at 1,024bp downstream of the end of *Asz1* exon 2. The HDR template was a double-stranded DNA fragment containing two loxP sites flanking exon 1 and exon 2 of *Asz1*. To generate *Asz1^cKO^* mice, *Stra8*-Cre transgenic mice (017490, Jackson Laboratory) were bred with *Asz1^flox/flox^* mice using the strategy as previously described. Primers used for *Asz1^flox^* genotyping PCR were 5’- GATAGCAGCATTGTATTCAAGG-3’ and 5’-CTAACCCCTAGTACTTCATGCTCC-3’. Primers used for *Stra8*-Cre genotyping PCR were 5’-GTGCAAGCTGAACAACAGGA-3’ and 5’-AGGGACACAGCATTGGAGTC-3’. Primers used for internal control in *Stra8*-Cre genotyping PCR were 5’-CTAGGCCACAGAATTGAAAGATCT-3’ and 5’- GTAGGTGGAAATTCTAGCATCATCC-3’.

For the generation of *Asz1^KO^* mice, *Stra8*-Cre *Asz1^flox/+^* mice were first crossed with WT C57BL/6 to obtain ASZ1*^+/-^* mice. *Asz1^KO^* mice were then generated by breeding *Asz1^+/-^* males and females. Primers used for *Asz1^KO^*genotyping PCR were 5’- TGGGTCTGACCTTCAACTGC-3’, 5’- TCCTCTCTCCTCTTCCGTGG-3’ and 5’- CTAACCCCTAGTACTTCATGCTCC-3’.

*Miwi^KO^*, *Tdrkh^cKO^*, and *Mov10l1^cKO^*mice were generated and genotyped as previously described^28, 55^.

### Antibody generation

For the generation of antibodies against TDRD7, DNA fragment encoding amino acids 1-246 of TDRD7 was cloned into pET-28a (His-tag) vectors. The resulting His-tagged recombinant proteins were expressed in *E. coli,* affinity purified and used as the antigen to generate rabbit polyclonal antisera (Pacific Immunology). The antisera were affinity-purified with the corresponding His-tagged antigens using AminoLink Plus immobilization kit (44894, Thermo Scientific).

### Histology

Testes and epididymides from adult WT and mutant mice were collected and fixed in Bouin’s fixative (HT10132, Sigma-Aldrich) overnight at 4°C and embedded in paraffin. For the histological analysis, sections were cut at 5μm, dewaxed, rehydrated, and stained with hematoxylin and eosin.

### Plasmid construction

The plasmids GFP-MIWI, TDRKH-RFP and TDRKH-GFP were constructed in our previous study^28^. The full-length *Asz1* cDNA was amplified by PCR and cloned into the pEGFP-C1 (GFP-tag at N-terminus) expression vector. The full-length cDNAs for *Gpat2* and *Pld6* were amplified by PCR and cloned into the pEGFP-N1 (GFP-tag at C-terminus) expression vector. To obtain Flag-tagged ASZ1, MILI, and MOV10L1 plasmids, the full-length cDNAs for *Asz1*, *Mili*, and *Mov10l1* were amplified by PCR and cloned into the pcDNA3-Flag (Flag-tag at N-terminus) expression vector.

### HeLa cell transfection and immunofluorescence

For Mitotracker staining, HeLa cells were transfected with indicated plasmids using Lipofectamine 3000 (L3000015, Thermo Fisher Scientific) according to the manufacturer’s instructions. After 24 h, cells were incubated with 300 nM Mitotracker Red Probes (M7512, Thermo Scientific) at 37°C for 30 min. Cells were then fixed with 4% PFA for 15 min, incubated in PBS containing 0.2% Triton X-100 for 20 min, and washed with PBS. Cells were mounted using Vectorshield mounting media with DAPI (H1200, Vector Laboratories) and imaged with Fluoview FV1000 confocal microscope (Olympus, Japan).

For fixed cell immunofluorescence, HeLa cells were transfected with indicated plasmids using Lipofectamine 3000 (L3000015, Thermo Fisher Scientific) according to the manufacturer’s instructions. After 24 h, HeLa cells were fixed with 4% PFA for 15 min, incubated in 0.2% Triton X-100 in PBS for 20 min, and washed with PBS. HeLa cells were blocked in 5% NGS at RT for 30 min and then incubated with anti-FLAG antibody (1:100; F1804, Sigma-Aldrich) overnight at 4°C. After washing with PBS, HeLa cells were incubated with Alexa Fluor 555 goat anti-mouse IgG (1:200; A21422, Thermo Fisher Scientific) at RT for 1 h. Cells were mounted using Vectashield mounting media with DAPI (H-1200, Vector Laboratories) and imaged using Fluoview FV1000 confocal microscope (Olympus, Japan).

### Western blotting

Mouse testes were collected and homogenized in RIPA buffer (J63306-AP, Thermo Fisher Scientific) with protease inhibitors (A32965, Thermo Fisher Scientific). Protein lysates were separated by 4–20% polyacrylamide gels (4561096, Bio-Rad) and transferred to PVDF membranes (1620177, Bio-Rad). After blocking in 5% non-fat milk at RT for 30 min, the membranes were incubated with primary antibodies in 5% non-fat milk overnight at 4°C. The primary antibodies used were anti-ASZ1 (1:500; 21550-1-AP, Proteintech), anti-MIWI (1:1000; 2079, Cell Signaling Technology), anti-MILI (1:2000; PM044, MBL), anti-MOV10L1 (1:500; kindly provided by P. Jeremy Wang), anti-MVH (1:4000; Ab27591, Abcam), anti-TDRKH (1:4000; 13528-1-AP, Proteintech), anti-TDRD1^35^ (1:1000), anti-TDRD5^36^ (1:1000), anti-TDRD6^46^ (1:1000), anti-TDRD7 (1:1000; homemade), or HRP-conjugated mouse anti-β-actin (1:5000; A3854, Sigma-Aldrich). Membranes were washed with TBST for 3 times and incubated with HRP-conjugated goat anti-rabbit IgG (1:5000; 1706515, Bio-Rad) at RT for 1 h followed by chemiluminescent detection with the ECL Substrate (1705060, Bio-Rad).

### Immunoprecipitation of piRNAs

Mouse testes were collected and homogenized in lysis buffer (20 mM HEPES pH 7.3, 150 mM NaCl, 2.5 mM MgCl_2_, 0.3% NP-40, and 1 mM DTT) containing protease inhibitors (A32965, Thermo Fisher Scientific) and RNase inhibitor (N2615, Promega). Protein A agarose beads (11134515001, Sigma-Aldrich) were added to the precleared lysates at 4°C for 2 h. Anti-MIWI (2079, Cell Signaling Technology) or anti-MILI (PM044, MBL) antibody together with Protein A agarose beads were added to the lysates and incubated at 4°C for 4 h. Beads were washed in lysis buffer for 5 times, and immunoprecipitated RNAs were isolated from the beads using Trizol reagent (15596026, Thermo Fisher Scientific) for piRNA labeling or small RNA library construction. For protein detection, immunoprecipitated beads were boiled in protein loading buffer for 5 min flowed by Western blotting of MIWI or MILI.

### Detection of piRNAs

Total RNA was extracted from mouse testes using Trizol reagent (15596026, Thermo Fisher Scientific). Total RNA or immunoprecipitated RNA (MIWI- or MILI-associated) was dephosphorylated with Shrimp Alkaline Phosphatase (M0371, NEB). RNA end-labeling was carried out using T4 polynucleotide kinase (M0201, NEB) and [γ-^32^P] ATP (NEG002A250UC, PerkinElmer). The ^32^P-labeled RNA was separated by 15% Urea-PAGE gel, and radioactive signal was examined by exposing the gel on phosphorimager screen followed by scanning on the Typhoon scanner (GE Healthcare).

### Small RNA libraries and bioinformatics

Small RNA libraries were prepared from total RNA using Small RNA Library Prep Kit (E7300, NEB) according to the manufacturer’s instructions. Indexed libraries were pooled and sequenced with the Illumina HiSeq 2500 or NovaSeq 6000 platform (MSU Genomic Core Facility). Sequenced reads were processed with fastx_clipper (FASTX-Toolkit) to remove sequencing adapters. Clipped reads were filtered by length (24–32 nt) and aligned with Bowtie, allowing one base mismatch, to the following sets of sequences: piRNA clusters, coding RNA, non-coding RNA, repeats, intron, and other. Repeats were classified according to RepeatMasker. For total small RNA sequencing, RNA read counts were normalized based on miRNA counts (21-23 nt).

### Immunoprecipitation and mass spectrometry (IP-MS)

Mouse testes were homogenized using lysis buffer (20 mM HEPES pH 7.3, 150 mM NaCl, 2.5 mM MgCl_2_, 0.3% NP-40, and 1 mM DTT) with protease inhibitors (04693132001, Sigma-Aldrich). The lysates were pre-cleared with Protein A agarose beads (11134515001, Sigma-Aldrich) at 4°C for 2 h, incubated with Rabbit IgG (2729S, Cell Signaling Technology) or anti-ASZ1 (21550-1-AP, Proteintech) antibody at 4°C overnight, followed by Protein A agarose beads at 4°C for 3 h. Beads were washed with lysis buffer and PBS, and proteins were eluted with 2% SDS at 90°C for 10 min. Eluates were digested with trypsin (T1426, Sigma-Aldrich) using the SP3 protocol^56^.

Peptides were resuspended in 5 μl of 2% acetonitrile/0.1% trifluoroacetic acid in water and separated on Thermo Acclaim PepMap RSLC C18 column using a 5%-24%-38%-90% 35-min gradient (mobile phase A, 0.1% formic acid in water; mobile phase B, 80% acetonitrile/0.1% formic acid in water). Eluted peptides were sprayed via a FlexSpray source into a Thermo Q-Exactive HF-X mass spectrometer, acquiring survey scans at 60,000 resolution (m/z 200) and fragmenting the top ten ions by HCD with MS/MS at 15,000 resolution.

The resulting MS/MS spectra were converted to peak lists using Mascot Distiller and searched against a Mus musculus protein reference database containing all sequences available from Uniprot appended with common laboratory contaminants using the Mascot searching algorithm^57^. The Mascot output was then analyzed using Scaffold to probabilistically validate protein identifications. Assignments validated using the Scaffold 1% FDR confidence filter were considered true.

### Transmission electron microscopy

Mouse testes were fixed with 2.5% glutaraldehyde in 0.1 M cacodylate buffer at 4°C overnight. After washing with 0.1 M cacodylate buffer, the testes were post-fixed with 1% osmium tetroxide in 0.1 M cacodylate buffer at RT for 2 h. The tissues were then dehydrated through a graded ethanol series, infiltrated and embedded in Spurr’s resin. Ultrathin sections (70 nm) were stained with uranyl acetate and lead citrate. Images were acquired using JEOL 1400 Flash Transmission Electron Microscope (Japan Electron Optics Laboratory, Japan).

### Testis immunofluorescence

Mouse testes were fixed in 4% paraformaldehyde (PFA) in PBS at 4°C overnight and embedded in paraffin. Testis sections were cut at 5 μm, dewaxed and rehydrated. Antigen retrieval was carried out in Tris-EDTA buffer (pH 9.0) or sodium citrate buffer (pH 6.0). Testis sections were blocked in 5% normal goat serum (NGS) at RT for 30 min, then incubated with primary antibodies diluted in 5% NGS at 4°C overnight. Antibodies used were: anti-MIWI (1:100; 2079, Cell Signaling Technology), anti-MILI (1:100; PM044, MBL), anti-TDRKH (1:100; 13528-1-AP, Proteintech), anti-ACRV1 (1:50; 14040-1-AP, Proteintech), anti-ASZ1 (1:50; 21550-1-AP, Proteintech), anti-TDRD1^35^ (1:100), anti-TDRD5^36^ (1:100), anti-TDRD6^46^ (1:100), anti-TDRD7 (1:100; homemade), anti-MOV10L1^58^ (1:100), anti-DDX25 (1:50; sc-166289, Santa Cruz Biotechnology), anti-LINE1 ORF1^59^ (1:800), or FITC-conjugated mouse anti-γH2AX (1:500; 16-202A, Millipore). After washing with PBS, sections were incubated with secondary antibodies at RT for 1 h and mounted using Vectashield mounting media with DAPI (H-1200, Vector Laboratories). Secondary antibodies used were Alexa Fluor 555 goat anti-rabbit IgG (1:500; A21429, Thermo Fisher Scientific), Alexa Fluor 488 goat anti-mouse IgG (1:500; A11029, Thermo Fisher Scientific). Fluorescence images were captured using Fluoview FV1000 confocal microscope (Olympus, Japan).

For co-immunostaining of TDRKH or DDX25 with various piRNA factors, anti-TDRKH (1:100; AF6286, R&D) and anti-DDX25 (1:50; sc-166289, Santa Cruz Biotechnology) were used. For co-immunostaining of ASZ1 with various piRNA factors, testis sections were first incubated with MIWI, MILI, TDRD1, TDRD5, TDRD6, or TDRD7 antibodies, followed by the secondary antibody Alexa Fluor 555 anti-rabbit IgG. The sections were then re-blocked with 5% NGS and incubated with anti-ASZ1 (21550-1-AP, Proteintech) prelabeled with the Zenon Alexa Fluor 488 Rabbit IgG Labeling Kit (Z25302, Thermo Fisher Scientific) following manufacturer’s instructions. Fiji (ImageJ) plugin ComDet v.0.5.5 was used for quantitative colocalization analysis as previously described^60^.

### Quantitative RT-PCR

Total RNA was extracted from testes using TRIzol reagent and treated with TURBO DNA-free Kit (AM1907, Thermo Scientific). 1 μg of RNA was reverse transcribed with iScript cDNA Synthesis Kit (Bio-Rad). cDNA was diluted 4-fold and qPCR was performed using iTaq Universal SYBR Green Supermix (1725121, Bio-Rad) in the QuantStudio 5 Real-Time PCR System (Thermo Scientific). Three biological replications were performed. GAPDH was used as a reference gene.

Primers used for qPCR were as follows: pri-let7g forward (5’-3’)- GTACGGTGTGGACCTCATCA, pri-let7g reverse (5’-3’)- TCTTGCTGTGTCCAGGAAAG; pach44-2 forward (5’-3’)- AGTCTGTGTAGTAGTTTCCTGAG, pach44-2 reverse (5’-3’)- TGTCCACTTCCATGTTACCT; pach39-1 forward (5’-3’)-GTTGCCCCAAGAGAATGTGT, pach39-1 reverse (5’-3’)- TTCCACAGGTCCAGCCTTAG; pach78-2 forward (5’-3’)- GTGAAGCTAAGGATGCTGGGATAG, pach78-2 reverse (5’-3’)- ACAGGATGTCCCCTGAAATCAGTC; pach78-1 forward (5’-3’)- TGTTCACTTACATATCAGGGTC, pach78-1 reverse (5’-3’)- GTAAAGCCCAAGAGCAAGAG; pach84-1 forward (5’-3’)- CTCCCAATGGCAACATCTTT, pach84-1 reverse (5’-3’)-TGACCCTTCAGGGAATTCAG; pach54 forward (5’-3’)-GATAACCACCAGCAGTTCTCCAC, pach54 reverse (5’-3’)- ACTCCTCATTGGTCCTTGTCTTG; pach98 forward (5’-3’)- GTTAGCGAAGGACATTATTCTAACC, pach98 reverse (5’-3’)- TGACATGAACACAGGTGCTCAGAT; pach44-1 forward (5’-3’)- ATTTCCTGGCTGGATGGTTGT, pach44-1 reverse (5’-3’)- AGGAGTGATGTGAGCGAGTGC; pach67 forward (5’-3’)- CTATGCTTATGATGGCATTGGAGAG, pach67 reverse (5’-3’)- TTCCAGTTCAACAGGGACACGGGAC; GAPDH forward (5’-3’)- AGAAACCTGCCAAGTATGATGAC, GAPDH reverse (5’-3’)- GTCATTGAGAGCAATGCCAG.

### Statistical analysis

The number of biological replicates (n) is indicated in each figure legend. Data are presented as mean ± SEM. Statistical comparisons between groups were performed using unpaired t-test, with significance defined as **P < 0.01.

## Data availability

All sequencing data have been deposited in the Sequence Read Archive of NCBI under the accession number PRJNA1347438. Data for IP-MS have been deposited at ProteomeXchange under the accession number PXD069037.

## Acknowledgements

We thank X. Cheng for critical reading of the manuscript. We thank P. J. Wang for MOV10L1 antibody. We thank D. Whitten for assistance in mass spectrometry. This work was supported by grants from National Institute of General Medical Sciences R35GM156209 (CC) and R01GM132490 (CC), Eunice Kennedy Shriver National Institute of Child Health and Human Development R01HD084494 (CC), and National Institute of Food and Agriculture MICL02690 (CC).

## Author contributions

X.Y., C.W., and C.C. designed research. X.Y., C.W., J.M.M., and Q.W. performed the research. H.X. and E.Y.D. generated *ASZ1* flox mice. L.S., G.S., and D.D. analyzed data. X.Y., C.W., and C.C. wrote the manuscript. C.C. supervised the project.

## Supplementary information

**Fig. S1.**
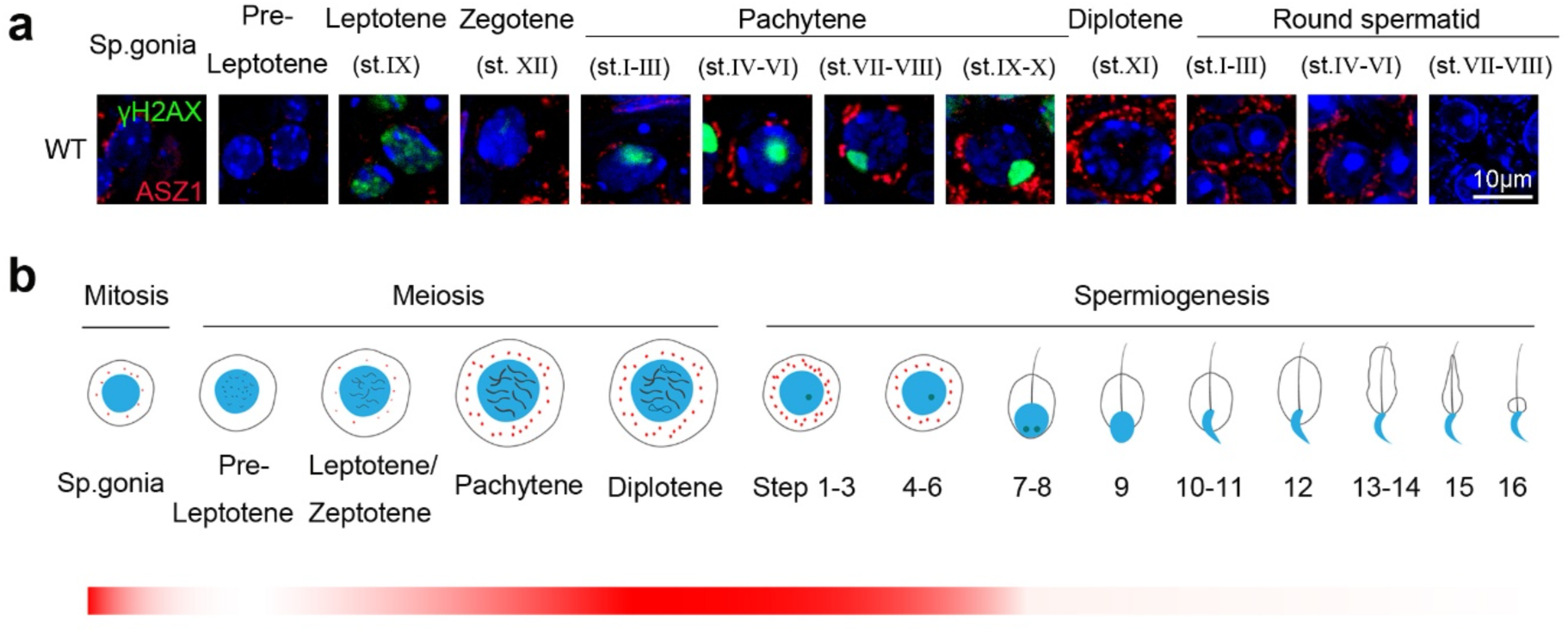
The expression pattern of ASZ1 during mouse spermatogenesis. **a** Immunofluorescence of ASZ1 and γH2AX in adult WT testes. **b** A schematic summary of the localization of ASZ1 in adult WT testes during spermatogenesis.

**Fig. S2.**
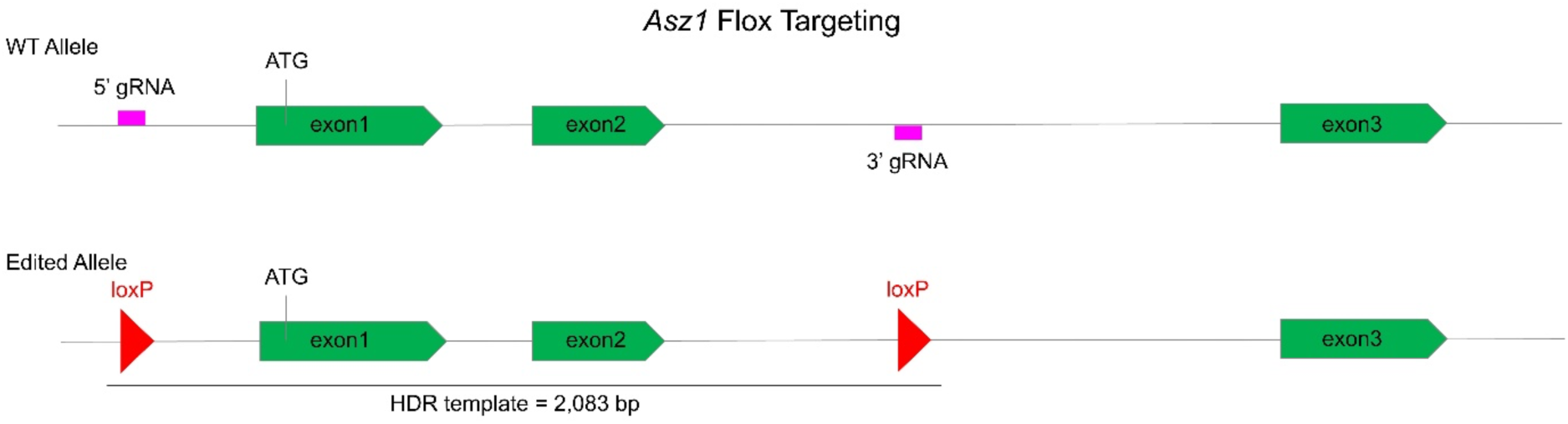
Targeting of the *Asz1* locus to generate *Asz1* flox mice. The location of gRNAs is indicated on the WT allele. The homology-directed repair (HDR) template containing two loxP sequences is indicated on the edited allele.

**Fig. S3.**
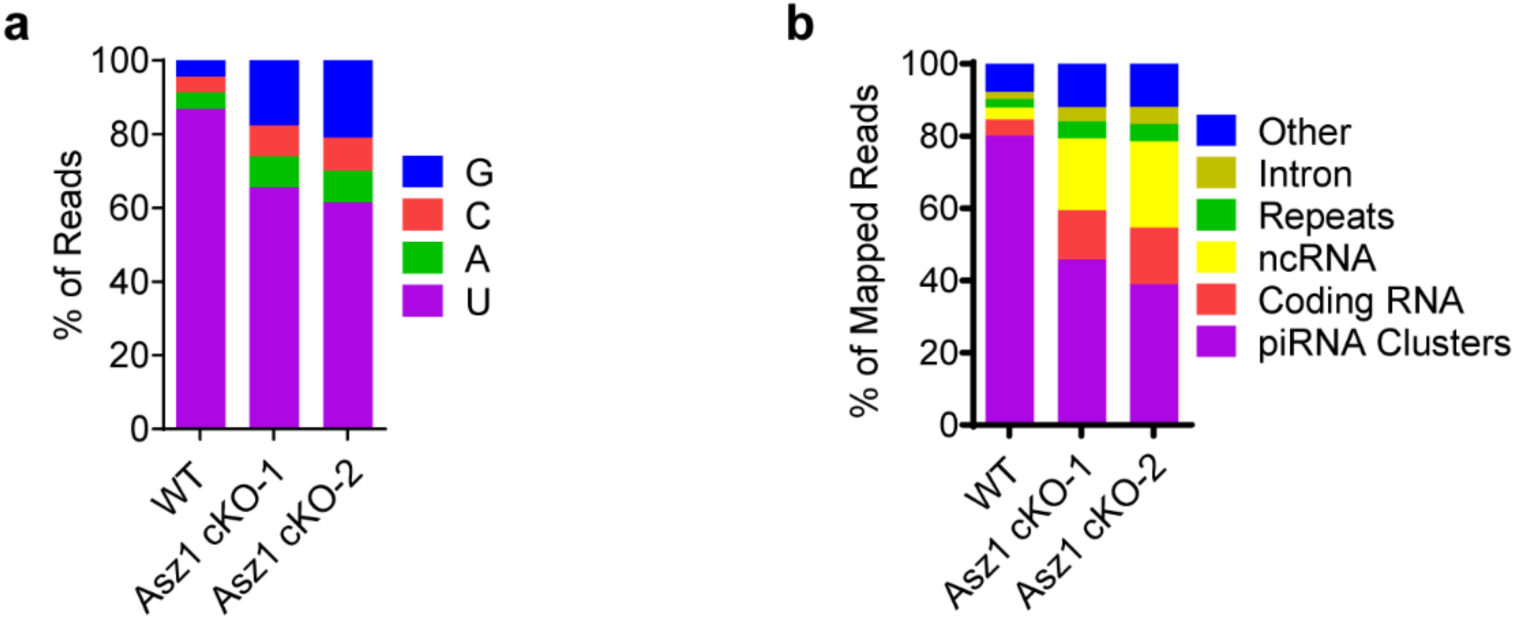
Primary piRNA pathway is severely disrupted in *Asz1* cKO testes. **a** Nucleotide distributions at the first position from adult WT and *Asz1^cKO^* testicular total small RNA libraries. **b** Genomic annotation of total piRNAs from adult WT and *Asz1^cKO^* testes. Sequence reads (24-32 nt) from testicular total small RNA libraries were aligned to mouse genomic sequence sets in the following order: piRNA clusters, coding RNA, non-coding RNA, repeats, intron, and other.

**Fig. S4.**
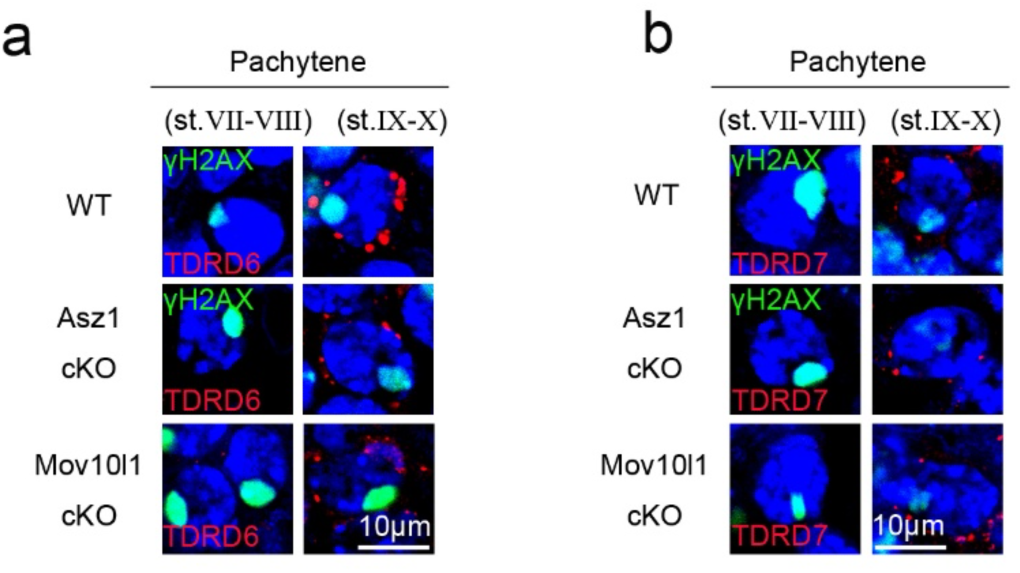
Immunostaining of TDRD6 and TDRD7 in WT, *Asz1^cKO^*, and *Mov10l1^cKO^* testes. **a** Immunofluorescence of TDRD6 and γH2AX in adult testis sections from WT, *Asz1^cKO^*, and *Mov10l1^cKO^* mice. **b** Immunofluorescence of TDRD7 and γH2AX in adult testis sections from WT, *Asz1^cKO^*, and *Mov10l1^cKO^* mice.

**Fig. S5.**
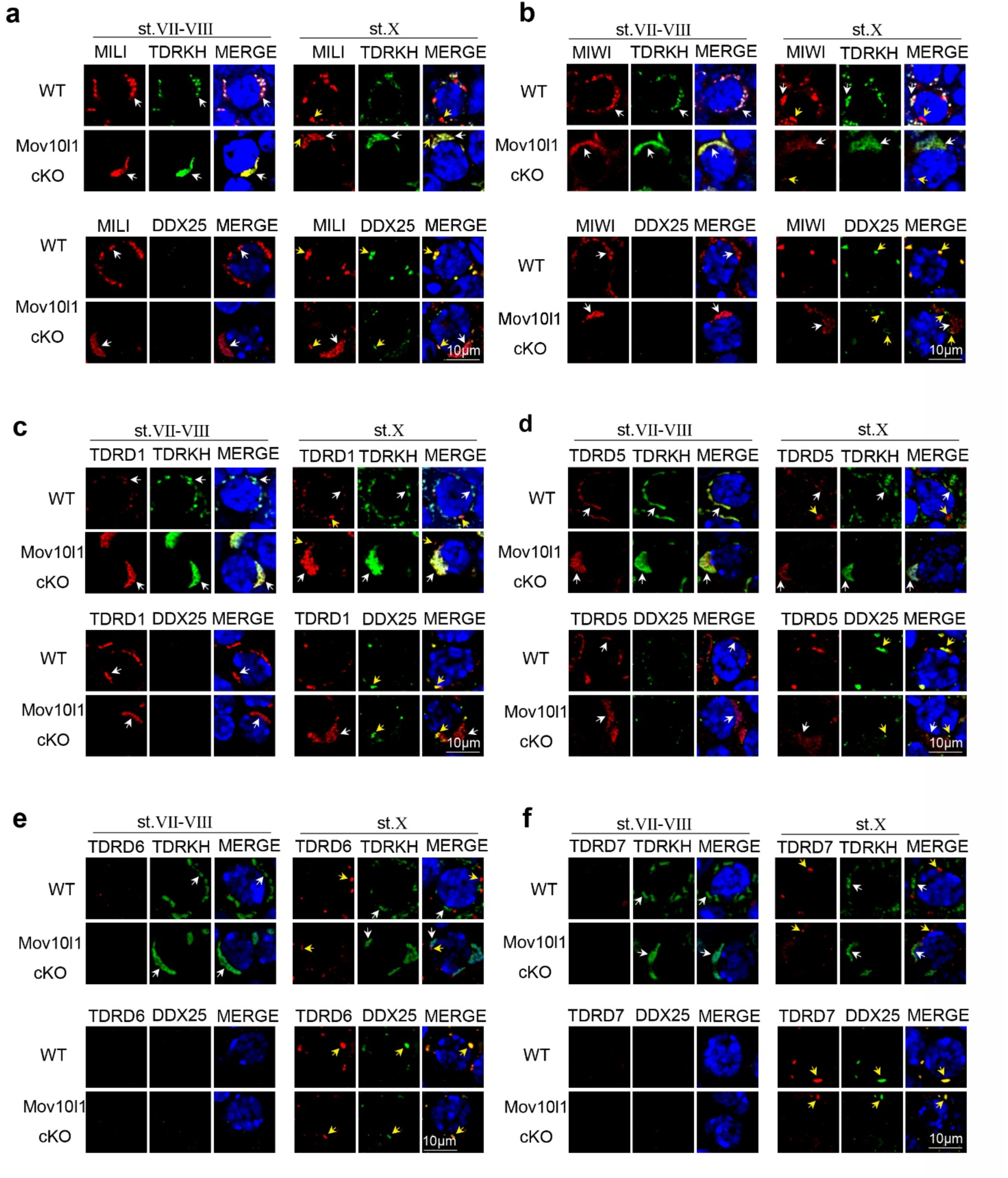
Loss of MOV10L1 impedes the translocation of piRNA pathway proteins from the IMC to CB precursors. **a** Co-immunostaining of MILI with TDRKH or DDX25 in adult testis sections from WT and *Mov10l1^cKO^* mice. **b** Co-immunostaining of MIWI with TDRKH or DDX25 in adult WT and *Mov10l1^cKO^* testes. **c** Co-immunostaining of TDRD1 with TDRKH or DDX25 in adult WT and *Mov10l1^cKO^* testes. **d** Co-immunostaining of TDRD5 with TDRKH or DDX25 in adult WT and *Mov10l1^cKO^* testes. **e** Co-immunostaining of TDRD6 with TDRKH or DDX25 in adult WT and *Mov10l1^cKO^* testes. **f** Co-immunostaining of TDRD7 with TDRKH or DDX25 in adult WT and *Mov10l1^cKO^* testes. White arrows indicate the IMC. Yellow arrows indicate CB precursors. All immunostaining images of WT are identical to those shown for WT in Fig. 7.

## References

1. Czech, B. et al. piRNA-Guided Genome Defense: From Biogenesis to Silencing. Annu Rev Genet 52, 131–157 (2018).

2. Ozata, D.M., Gainetdinov, I., Zoch, A., O’Carroll, D. & Zamore, P.D. PIWI-interacting RNAs: small RNAs with big functions. Nat Rev Genet 20, 89–108 (2019).

3. Wang, X., Ramat, A., Simonelig, M. & Liu, M.F. Emerging roles and functional mechanisms of PIWI-interacting RNAs. Nat Rev Mol Cell Biol (2022).

4. Siomi, M.C., Sato, K., Pezic, D. & Aravin, A.A. PIWI-interacting small RNAs: the vanguard of genome defence. Nat Rev Mol Cell Biol 12, 246–258 (2011).

5. Thomson, T. & Lin, H. The biogenesis and function of PIWI proteins and piRNAs: progress and prospect. Annu Rev Cell Dev Biol 25, 355–376 (2009).

6. Gainetdinov, I., Colpan, C., Arif, A., Cecchini, K. & Zamore, P.D. A Single Mechanism of Biogenesis, Initiated and Directed by PIWI Proteins, Explains piRNA Production in Most Animals. Mol Cell 71, 775–790 e775 (2018).

7. Honda, S. et al. Mitochondrial protein BmPAPI modulates the length of mature piRNAs. RNA 19, 1405–1418 (2013).

8. Koga, Y. et al. Dipteran-specific Daedalus controls Zucchini endonucleolysis in piRNA biogenesis independent of exonucleases. Cell Rep 43, 114923 (2024).

9. Munafo, M. et al. Daedalus and Gasz recruit Armitage to mitochondria, bringing piRNA precursors to the biogenesis machinery. Genes Dev 33, 844–856 (2019).

10. Ge, D.T. et al. The RNA-Binding ATPase, Armitage, Couples piRNA Amplification in Nuage to Phased piRNA Production on Mitochondria. Mol Cell 74, 982–995 e986 (2019).

11. Watanabe, T. et al. MITOPLD is a mitochondrial protein essential for nuage formation and piRNA biogenesis in the mouse germline. Dev Cell 20, 364–375 (2011).

12. Li, X.Z. et al. An ancient transcription factor initiates the burst of piRNA production during early meiosis in mouse testes. Mol Cell 50, 67–81 (2013).

13. Girard, A., Sachidanandam, R., Hannon, G.J. & Carmell, M.A. A germline-specific class of small RNAs binds mammalian Piwi proteins. Nature 442, 199–202 (2006).

14. Aravin, A. et al. A novel class of small RNAs bind to MILI protein in mouse testes. Nature 442, 203–207 (2006).

15. Lau, N.C. et al. Characterization of the piRNA complex from rat testes. Science 313, 363–367 (2006).

16. Ozata, D.M. et al. Evolutionarily conserved pachytene piRNA loci are highly divergent among modern humans. Nat Ecol Evol 4, 156–168 (2020).

17. Aravin, A.A. Pachytene piRNAs as beneficial regulators or a defense system gone rogue. Nat Genet 52, 644–645 (2020).

18. Xu, M.M. & Li, X.Z. Enigmatic Pachytene PIWI-Interacting RNAs. Genome Biol Evol 16 (2024).

19. Wu, P.H. et al. The evolutionarily conserved piRNA-producing locus pi6 is required for male mouse fertility. Nat Genet 52, 728–739 (2020).

20. Choi, H., Wang, Z. & Dean, J. Sperm acrosome overgrowth and infertility in mice lacking chromosome 18 pachytene piRNA. PLoS Genet 17, e1009485 (2021).

21. Kuramochi-Miyagawa, S. et al. Mili, a mammalian member of piwi family gene, is essential for spermatogenesis. Development 131, 839–849 (2004).

22. Deng, W. & Lin, H. miwi, a murine homolog of piwi, encodes a cytoplasmic protein essential for spermatogenesis. Dev Cell 2, 819–830 (2002).

23. Aravin, A.A. et al. A piRNA pathway primed by individual transposons is linked to de novo DNA methylation in mice. Mol Cell 31, 785–799 (2008).

24. De Fazio, S. et al. The endonuclease activity of Mili fuels piRNA amplification that silences LINE1 elements. Nature 480, 259–263 (2011).

25. Reuter, M. et al. Miwi catalysis is required for piRNA amplification-independent LINE1 transposon silencing. Nature 480, 264–267 (2011).

26. Chen, C., Nott, T.J., Jin, J. & Pawson, T. Deciphering arginine methylation: Tudor tells the tale. Nat Rev Mol Cell Biol 12, 629–642 (2011).

27. Chen, C. et al. Mouse Piwi interactome identifies binding mechanism of Tdrkh Tudor domain to arginine methylated Miwi. Proc Natl Acad Sci U S A 106, 20336–20341 (2009).

28. Ding, D. et al. Mitochondrial membrane-based initial separation of MIWI and MILI functions during pachytene piRNA biogenesis. Nucleic Acids Res 47, 2594–2608 (2019).

29. Ma, L. et al. GASZ is essential for male meiosis and suppression of retrotransposon expression in the male germline. PLoS Genet 5, e1000635 (2009).

30. Saxe, J.P., Chen, M., Zhao, H. & Lin, H. Tdrkh is essential for spermatogenesis and participates in primary piRNA biogenesis in the germline. EMBO J 32, 1869–1885 (2013).

31. Shiromoto, Y. et al. GPAT2, a mitochondrial outer membrane protein, in piRNA biogenesis in germline stem cells. RNA 19, 803–810 (2013).

32. Huang, H. et al. piRNA-associated germline nuage formation and spermatogenesis require MitoPLD profusogenic mitochondrial-surface lipid signaling. Dev Cell 20, 376–387 (2011).

33. Vourekas, A. et al. The RNA helicase MOV10L1 binds piRNA precursors to initiate piRNA processing. Genes Dev 29, 617–629 (2015).

34. Zheng, K. & Wang, P.J. Blockade of pachytene piRNA biogenesis reveals a novel requirement for maintaining post-meiotic germline genome integrity. PLoS Genet 8, e1003038 (2012).

35. Gao, J. et al. TDRD1 phase separation drives intermitochondrial cement assembly to promote piRNA biogenesis and fertility. Dev Cell 59, 2704–2718 e2706 (2024).

36. Ding, D. et al. TDRD5 binds piRNA precursors and selectively enhances pachytene piRNA processing in mice. Nat Commun 9, 127 (2018).

37. Tanaka, T. et al. Tudor domain containing 7 (Tdrd7) is essential for dynamic ribonucleoprotein (RNP) remodeling of chromatoid bodies during spermatogenesis. Proc Natl Acad Sci U S A 108, 10579–10584 (2011).

38. Castaneda, J. et al. Reduced pachytene piRNAs and translation underlie spermiogenic arrest in Maelstrom mutant mice. EMBO J 33, 1999–2019 (2014).

39. Di Giacomo, M. et al. Multiple epigenetic mechanisms and the piRNA pathway enforce LINE1 silencing during adult spermatogenesis. Mol Cell 50, 601–608 (2013).

40. Wei, C. et al. PNLDC1 catalysis and postnatal germline function are required for piRNA trimming, LINE1 silencing, and spermatogenesis in mice. PLoS Genet 20, e1011429 (2024).

41. Ipsaro, J.J., Haase, A.D., Knott, S.R., Joshua-Tor, L. & Hannon, G.J. The structural biochemistry of Zucchini implicates it as a nuclease in piRNA biogenesis. Nature 491, 279–283 (2012).

42. Nishimasu, H. et al. Structure and function of Zucchini endoribonuclease in piRNA biogenesis. Nature 491, 284–287 (2012).

43. Chuma, S. et al. Tdrd1/Mtr-1, a tudor-related gene, is essential for male germ-cell differentiation and nuage/germinal granule formation in mice. Proc Natl Acad Sci U S A 103, 15894–15899 (2006).

44. Yabuta, Y. et al. TDRD5 is required for retrotransposon silencing, chromatoid body assembly, and spermiogenesis in mice. J Cell Biol 192, 781–795 (2011).

45. Vasileva, A., Tiedau, D., Firooznia, A., Muller-Reichert, T. & Jessberger, R. Tdrd6 is required for spermiogenesis, chromatoid body architecture, and regulation of miRNA expression. Curr Biol 19, 630–639 (2009).

46. Wei, H. et al. piRNA loading triggers MIWI translocation from the intermitochondrial cement to chromatoid body during mouse spermatogenesis. Nat Commun 15, 2343 (2024).

47. Lehtiniemi, T. & Kotaja, N. Germ granule-mediated RNA regulation in male germ cells. Reproduction 155, R77–R91 (2018).

48. Gao, J. et al. PIWI proteins tether the piRNA biogenesis machinery to mitochondria during mammalian spermatogenesis. EMBO J (2025).

49. Miao, J. et al. GASZ directly recruits MILI to the intermitochondrial cement for piRNA biogenesis and male germ cell development. Nucleic Acids Res 53 (2025).

50. Wasik, K.A. et al. RNF17 blocks promiscuous activity of PIWI proteins in mouse testes. Genes Dev 29, 1403–1415 (2015).

51. Gou, L.T. et al. Pachytene piRNAs instruct massive mRNA elimination during late spermiogenesis. Cell Res 24, 680–700 (2014).

52. Dai, P. et al. A Translation-Activating Function of MIWI/piRNA during Mouse Spermiogenesis. Cell 179, 1566–1581 e1516 (2019).

53. Cecchini, K., Ajaykumar, N., Bagci, A., Zamore, P.D. & Gainetdinov, I. Mouse Pachytene piRNAs Cleave Hundreds of Transcripts, But Alter the Steady-State Abundance of Only a Minority of Targets. bioRxiv (2024).

54. Vourekas, A. et al. Mili and Miwi target RNA repertoire reveals piRNA biogenesis and function of Miwi in spermiogenesis. Nat Struct Mol Biol 19, 773–781 (2012).

55. Wei, C. et al. MIWI N-terminal RG motif promotes efficient pachytene piRNA production and spermatogenesis independent of LINE1 transposon silencing. PLoS Genet 19, e1011031 (2023).

56. Hughes, C.S. et al. Single-pot, solid-phase-enhanced sample preparation for proteomics experiments. Nature protocols 14, 68–85 (2019).

57. Perkins, D.N., Pappin, D.J., Creasy, D.M. & Cottrell, J.S. Probability-based protein identification by searching sequence databases using mass spectrometry data. ELECTROPHORESIS: An International Journal 20, 3551–3567 (1999).

58. Zheng, K. et al. Mouse MOV10L1 associates with Piwi proteins and is an essential component of the Piwi-interacting RNA (piRNA) pathway. Proc Natl Acad Sci U S A 107, 11841–11846 (2010).

59. Ding, D. et al. PNLDC1 is essential for piRNA 3’ end trimming and transposon silencing during spermatogenesis in mice. Nat Commun 8, 819 (2017).

60. Chung, P.Y., Shoji, K., Izumi, N. & Tomari, Y. Dynamic subcellular compartmentalization ensures fidelity of piRNA biogenesis in silkworms. EMBO reports 22, e51342 (2021)

